# Immune cells employ intermittent integrin-mediated traction forces for 3D migration

**DOI:** 10.1101/2023.04.20.537658

**Authors:** Tina Czerwinski, Lars Bischof, David Böhringer, Sibel Kara, Pamela L. Strissel, Reiner Strick, Alexander Winterl, Richard Gerum, Ernst Wittmann, Michael Schneider, Matthias W. Beckmann, Gina Nusser, Manuel Wiesinger, Silvia Budday, Anja Lux, Caroline Voskens, Ben Fabry, Christoph Mark

## Abstract

To reach targets outside the bloodstream, immune cells can extravasate and migrate through connective tissue. During tissue infiltration, immune cells migrate in an amoeboid fashion, characterized by weak matrix adhesions and low traction forces, that allows them to achieve high migration speeds of up to 10 *µ*m/min. How immune cells reconcile amoeboid migration with the need to overcome steric hindrance in dense matrices is currently not understood. Here we show that NK92 (natural killer) cells can switch from their default amoeboid migration mode to a contractile, mesenchymal-like migration mode when moving through fibrous human amniotic membrane (HAM) tissue. We subsequently study immune cell migration in reconstituted 3D collagen networks with known mechanical properties and pore sizes and apply time-lapse confocal reflection microscopy to obtain simultaneous measurements of migration speed, directional persistence, and cell contractility. We find that NK92 cells are highly mechanoresponsive and exert substantial acto-myosin driven, integrin-mediated contractile forces of up to 100 nN on the extracellular matrix during short contractile phases. This burst-like contractile behavior is also found in primary B, T, NK cells, neutrophils, and monocytes, and is tightly related to the fraction of cells that appear to become stuck in narrow pores of the surrounding matrix. Our results demonstrate that steric hindrance guides the rapid regulation of integrin-mediated adhesion to the ECM in a large number of immune cell subtypes.

## Introduction

Immune cells such as monocytes, neutrophilic granulocytes, as well as B cells, T cells, and natural killer (NK) cells can migrate through tissues up to 100 times faster than mesenchymal cells^1,2^. Immune cells typically use an amoeboid migration mode with limited or no proteolytic activity and limited or no specific (e.g. integrin-mediated) adhesion to the extracellular matrix (ECM)^3–5^. Therefore, these cells are not expected to generate substantial traction forces, implying that they do not form mature focal adhesion contacts and thus are not able to significantly pull on and rearrange ECM fibers^4,6^. However, since migration in three-dimensional (3D) environments requires the cell to overcome the resistive forces of the matrix^7–9^, some mechanism of force transmission across the plasma membrane to the ECM must exist but has not yet been characterized in these cells.

The involvement of integrins in leukocyte migration in 3D matrices has been studied previously and the general conclusion is that integrins are required during extravasation, but are not essential during tissue infiltration. For example, when integrin function is blocked in T cells or when integrins are depleted in mouse dendritic cells, neutrophils, and B cells, migration is only possible in confined 3D environments but not on two-dimensional (2D) surfaces^2,4^. More specifically, integrin depletion in mouse leukocytes does not affect migration speed in 3D collagen gels, indicating that these cells do not require integrins for interstitial migration, whereas they do for migration on 2D substrates^2^. Dendritic cells can switch from integrin-mediated to integrin-independent movement without any change in migration speed or directional persistence^10^. Furthermore, integrin-mediated force coupling is not required for migration in constrained environments. The currently established notion is that integrins in immune cells act as a low-affinity frictional interface between cortical actin flow and the substrate, thereby facilitating cell locomotion^11–13^.

In this study, we revisit this well-established notion, which, as our data on large traction forces generated by immune cells suggest, is not wrong but needs to be extended. We use human amniotic membrane (HAM) tissue as a tangible physiological model of interstitial tissue and track movements of NK92 cells using time-lapse confocal microscopy (Supplementary Videos 1-3). In addition, we monitor tissue deformations around migrating NK92 cells using confocal reflection microscopy. Consistent with prevailing theories, our data confirm that immune cells appear to migrate in an ameboid manner within fibrous, collagen-rich HAM tissue most of the time, without significantly adhering to or pulling on their micro-environment. Interestingly and unexpectedly, this amoeboid migration mode is frequently interrupted by short phases during which the NK92 cells induce substantial and long-ranging matrix deformations along their axis of movement.

Since the local mechanical properties of HAM tissue around a migrating cell are difficult to measure, we further perform migration experiments using reconstituted collagen gels with well-defined mechanical and structural properties. This allows us to apply fast time-lapse 3D traction force microscopy to reconstruct cell-generated contractile forces from the measured 3D deformations. We combine this technique with high-throughput 3D migration assays to relate cellular force generation to the migratory ability of immune cells under steric hindrance.

For NK92 cells embedded in 3D reconstituted collagen gels, we find that amoeboid migration is frequently interrupted by short contractile bursts (similar to our findings in amniotic tissue), with peak forces of up to ∼60 nN. In the following, we test the hypothesis that during force bursts, immune cells switch to a highly contractile mesenchymal-like migration mode that allows them to overcome the steric hindrance imposed by narrow pores in the ECM and to avoid getting “stuck” in the matrix (Supplementary Videos 4,5). We provide evidence for this hypothesis in three ways: first, we find a significant temporal correlation between cell contractility, cell speed, and directional persistence of individual motile immune cells. Second, we find that by blocking integrin adhesion to the matrix, NK92 cells can still migrate in an amoeboid migration mode with unchanged migration speed, but the cells are forced to take more frequent turns to avoid small pores and obstacles, and become “stuck” more frequently (reduced motile fraction). Third, we find that increasing the contractile forces of NK92 cells substantially increases the number of motile cells.

## Results

### Contractile phases of NK92 cells during migration in human amniotic membrane tissue

The human amniotic membrane (HAM) makes up the inner layer of the placenta and can be dissected while remaining intact for further use (Fig. 1a). HAM is widely used as a scaffold for tissue engineering in regenerative medicine^14^, or as a matrix to study cancer cell invasion^14,15^. With a shear modulus of 100-400 Pa, depending on preparation, HAM represents soft human tissue, which makes it an ideal system to detect cell-induced matrix deformations^16^. HAM consists of the amniotic surface layer containing epithelial polygonal cells, a basement membrane, and a fibrous extracellular matrix layer enriched in proteins including collagen I, collagen III, collagen IV, laminin, and fibronectin (Fig. 1b-d)^17^. Confocal reflection microscopy reveals a highly heterogeneous, fibrous structure of the ECM layer with a high variability in pore size (Fig. 1e). To test the ability of immune cells to invade such a mechanically challenging environment, we prepare HAM samples with the epithelial layer flipped downwards, and seed NK92 cells on top of the ECM layer. By employing time-lapse confocal imaging, we can track the movements of individual NK92 cells within the matrix, and at the same time track any deformations that the cells exert onto the matrix during 3D migration (Fig. 1f-i, Supplementary Videos 1-3).

**Figure 1:**
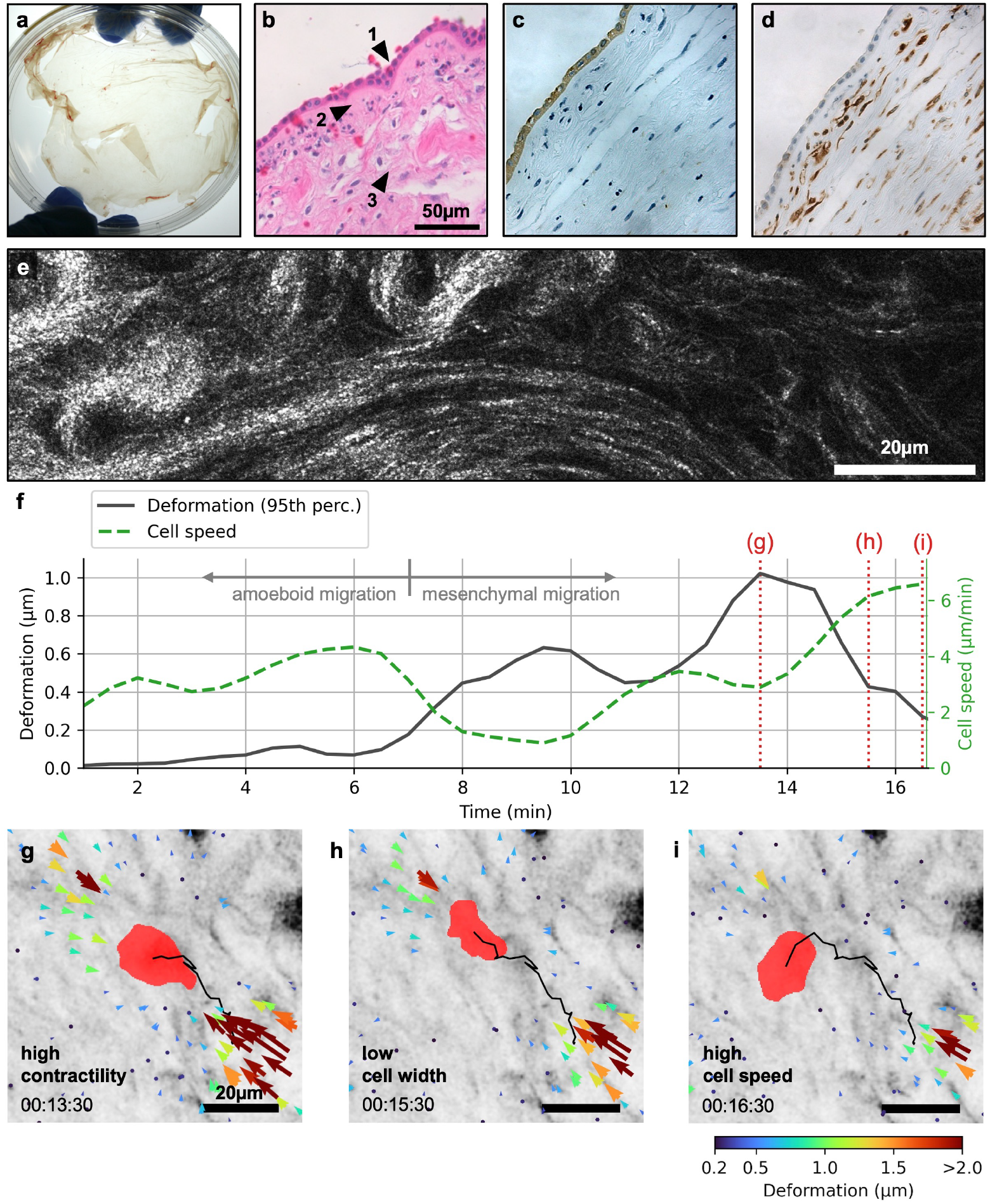
Migration of NK92 cells in ex-vivo amniotic membrane tissue. **a:** Macroscopic image of human amniotic membrane (HAM) tissue after fractionation from the placenta. **b:** Cross-sectional H&E stain of a HAM sample, showing the epithelial layer (1), the basement membrane (2), and the fibrous extracellular matrix layer (3). **c:** Cross-sectional pan-cytokeratin staining of a HAM sample, highlighting the epithelial layer (brown). **d:** Cross-sectional vimentin staining of a HAM sample, indicating the presence of fibroblasts in the fibrous extracellular matrix layer (brown). **e:** Exemplary confocal reflection image of a HAM sample showing the fibrous extracellular matrix layer and illustrating its structural heterogeneity, from thick, dense fiber bundles and small pore sizes on the left to thinner fibrils and larger pore sizes on the right. **f:** Time course of the matrix deformations (black line) and migration speed (green dashed line) of an exemplary NK92 cell migrating in HAM tissue. We use the 95^th^ percentile of all deformations within 100*µ*m around the cell to quantify highly localized deformations corresponding to contractile cell forces. Dotted red lines indicate the time steps corresponding to the deformation fields shown in panels (g), (h), and (i). **g:** Inverted collagen reflection image (gray) of an exemplary NK92 cell (red) migrating through HAM tissue. Arrows indicate strong contractile matrix deformations along the direction of cell movement (black line; c.f. panel f). **h:** Same as in (g), but two minutes later, when the cell has reduced its width perpendicular to the direction of movement (presumably to pass through a narrow pore), while matrix deformations are partially relaxed (c.f. panel f). **i:** Same as in (h), but one minute later, when matrix deformations are almost completely relaxed and the cell has attained a high migration speed again.

NK92 cells readily invade into the HAM tissue. They migrate predominantly without noticeable matrix deformations, supporting an amoeboid migration mode, but they occasionally switch into a different migration mode in which we observe substantial matrix deformations towards the cell center, predominantly along the direction of cell movement (Fig. 1f,g, Supplementary Videos 1-3). Hence, the cells appear to pull on the matrix in short contractile bursts that last only a few minutes. These bursts are frequently accompanied by a decrease in cell width, likely due to the cell moving through a narrow pore of the fiber network (Fig. 1h). Shortly after a contractile event, we typically observe an increase in migration speed (Fig. 1i).

### Contractility and mechanosensitivity of NK92 cells

While HAM tissue serves as a physiologically relevant model for interstitial tissue, it does not allow us to translate the measured deformations exerted by a cell into contractile forces, as we cannot quantify the local mechanical properties of the highly heterogeneous HAM tissue. We overcome this limitation by seeding NK92 cells into reconstituted collagen gels with known mechanical and structural properties, and apply high-speed 3D traction force microscopy^18^.

With a shear modulus of ∼100 Pa at a concentration of 0.6 mg/ml, ∼300 Pa at 1.2 mg/ml, and ∼1200 Pa at 2.4 mg/ml, reconstituted collagen gels accurately recapitulate the stiffness of HAM tissue (Supplementary Fig. 1). If we approximate the Young’s modulus as E = G·2·(1+*v*) with a Poisson ratio of *v*=0.25, we obtain a stiffness range for the collagen gels between ∼250-3000Pa, which covers the microelasticity of brain (400 Pa), liver (1350 Pa), kidney (2600 Pa), and fat tissue (3000 Pa)^19^. Due to the highly non-linear strain-stiffening behavior of collagen, the matrix surrounding an actively pulling NK cell may locally stiffen by a factor of 5 (Supplementary Figure 2). This widens the stiffness range that is sensed by the embedded immune cells, which then includes the microelasticity of lung (6000 Pa) and muscle (12kPa) tissue, covering a wide range of tissue stiffness on the cellular scale^19^.

For 3D traction force microscopy (TFM), we quantify the 3D matrix deformations around individual NK92 cells using particle image velocimetry applied to confocal reflection microscopy image stacks recorded over 24 min at a time interval of 1 min (Fig. 2a,b,e). Maximum matrix deformations are on the order of 0.25-0.5 *µ*m and comparable to the deformations we observe in HAM tissue (Fig. 2f). Matrix deformations tend to decrease with increasing collagen concentration and hence with increasing matrix stiffness (Fig. 2f).

**Figure 2:**
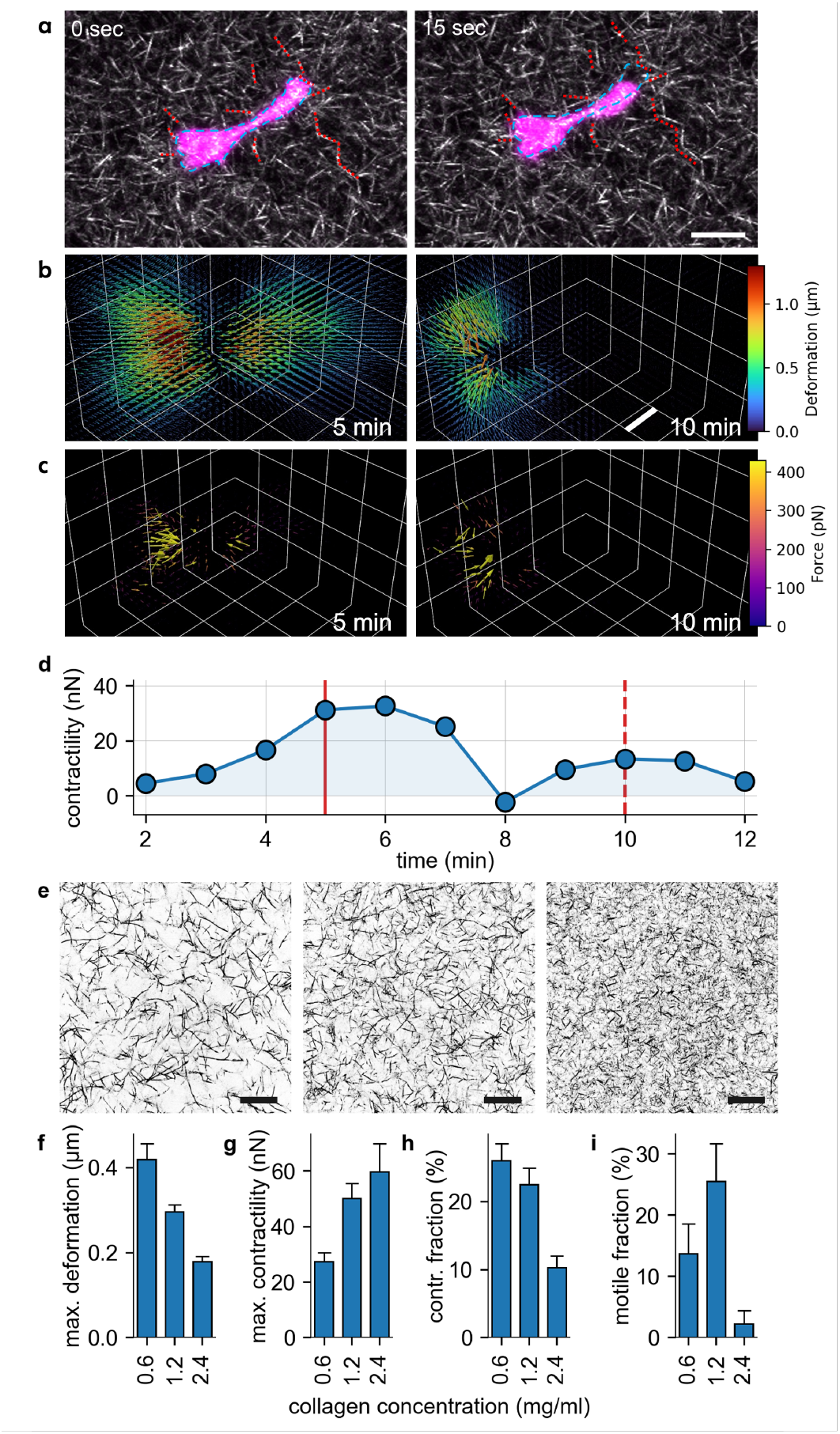
Force generation of NK92 cells in reconstituted collagen gels. **a:** Confocal image series of an ex-vivo expanded NK cell (magenta, Calcein stain) migrating in a 1.2 mg/ml collagen gel (imaged with confocal reflection microscopy). The dashed red lines indicate the position of exemplary collagen fibers at t=0s, to illustrate the movement of the fibers due to forces applied by the NK cell. The dashed blue line indicates the outline of the NK cell at t=0s. Scale bar: 10*µ*m. **b:** Measured 3D deformation field of the collagen fiber network that surrounds the NK cell at t=5min (corresponding to the red line in (d)) and at t=10min (corresponding to the dashed red line in (d)). Scale bar = 20*µ*m. **c:** Same as in (b), but showing the reconstructed force field that is obtained from the measured deformation fields. **d:** Temporal evolution of the contractility of a NK cell over the course of 10min. A typical contractile burst is visible during minutes 4-7. **e:** Confocal reflection image through a 3D reconstituted collagen gel with a collagen concentration of 0.6 mg/ml (shear modulus: 101 Pa; left), 1.2 mg/ml (286 Pa; center), and 2.4 mg/ml (1155 Pa; right). Collagen fibers are visible in black. Scale bar: 20 *µ*m. **f:** 99th percentile of the matrix deformations generated by NK92 cells for different collagen concentrations (0.6 mg/ml & 1.2 mg/ml: n=51 in 4 independent experiments; 2.4 mg/ml: n=46 in 3 independent experiments). **g:** Maximum cell contractility during a 24 min observation period. **h:** Fraction of contractile phases (time periods when cell contractility > 5 nN per cell). **i:** Fraction of motile NK92 cells for different collagen concentrations. A cell is defined as motile if the 5 min bounding-box around its migration path has a mean diagonal length of 6.5*µ*m or greater. Error bars denote 1 SEM.

We reconstruct the cell-generated forces responsible for the observed matrix deformations using high-speed 3D traction force microscopy (Fig. 2c), and subsequently sum up all inward pointing force vectors at each point in time to obtain the contractility of the cell over time (Fig. 2d)^8,18,20^. The method is based on a non-linear, non-affine material model for collagen that considers the force-induced strain stiffening and buckling of the collagen fibers^8,18^. As immune cells are expected to migrate without substantial force generation most of the time, we focus on the maximum contractility of each cell measured within the 24 min measurement period. We find that cell contractility increases with increasing matrix stiffness, reaching ∼60 nN at the highest collagen concentration (2.4 mg/ml, corresponding to 1155 Pa shear modulus; Fig. 2g; Supplementary Fig. 1). Given the small size of NK92 cells, this contractility is surprisingly high – comparable to mesenchymal cancer cells that generate forces of ∼100 nN^8,21^.

The increase in cell contractility with increasing collagen concentration further reveals that NK92 cells are able to sense mechanical changes of their environment and upregulate force generation in denser collagen gels, similar to the mechano-responsiveness seen in many mesenchymal cells^22,23^. However, denser collagen gels not only attain an increased gel stiffness but also a decreased pore size (Supplementary Fig. 1). We confirm that NK92 cells indeed adapt cell contractility specifically in response to matrix stiffness and not to pore size by repeating the experiment in a different collagen batch with approximately the same pore size but lower stiffness. We find approximately 2-fold lower contractile forces using the 4-fold softer batch (Supplementary Fig. 3).

In contrast to fibroblasts and cancer cells, NK92 cells are not always contractile but instead display short contractile bursts. We find that NK92 cells spend about 20-25% of the time in these contractile phases (> 5nN) when cultured in low– and medium-concentration collagen gels (Fig. 2h). In high-concentration collagen gels with narrower pores, the time spent in contractile phases drops to ∼10% (Fig. 2h).

To explain this decrease, we consider the motile phases of NK92 cells. We draw a bounding-box around a cell’s migration path during each 5 min period and determine its diagonal length (a measure of the exploration volume). A cell is classified as motile if the diagonal is greater than 6.5 *µ*m on average over the 24 min observation time. In low– and medium-concentration collagen gels, we find a significant fraction of motile cells, ranging between 10-30% (Fig. 2i). In high-concentration collagen gels, however, almost all cells remain non-motile (Fig. 2i). This behavior appears to scale with the product of maximum contractility and fraction of contractile phases (Fig. 2g-i). Accordingly, successful migration requires force bursts with sufficient magnitude and frequency. The dramatic decrease in the motile fraction in the high-concentration collagen gel suggests that narrow pores cause excessive steric hindrance beyond a level where cells cannot overcome necessary increases of magnitude or frequency of traction forces.

### Contractile phases correlate with increased cell speed and directional persistence

In high-concentration collagen gels, NK92 cells upregulate traction forces but cannot remain motile due to abundant narrow pores^24^. This leads to the question of whether traction forces can enhance motility in medium (and low) concentration collagen gels where pore size is not a limiting factor. Therefore, we investigate whether individual bursts of contractility lead to a higher cell speed and a directionally more persistent migration path (Fig. 3a,b). To this end, we track the position of individual cells during the 3D traction force microscopy measurement and extract the momentary cell speed and turning angle between subsequent time steps (see methods; Fig. 3a,b). As we are interested in the systematic movement of the cell as a whole, we employ Bayesian filtering to suppress spurious movements of the cell due to cell shape variations and imaging noise (see methods, Supplementary Figure 4)^25,26^.

**Figure 3:**
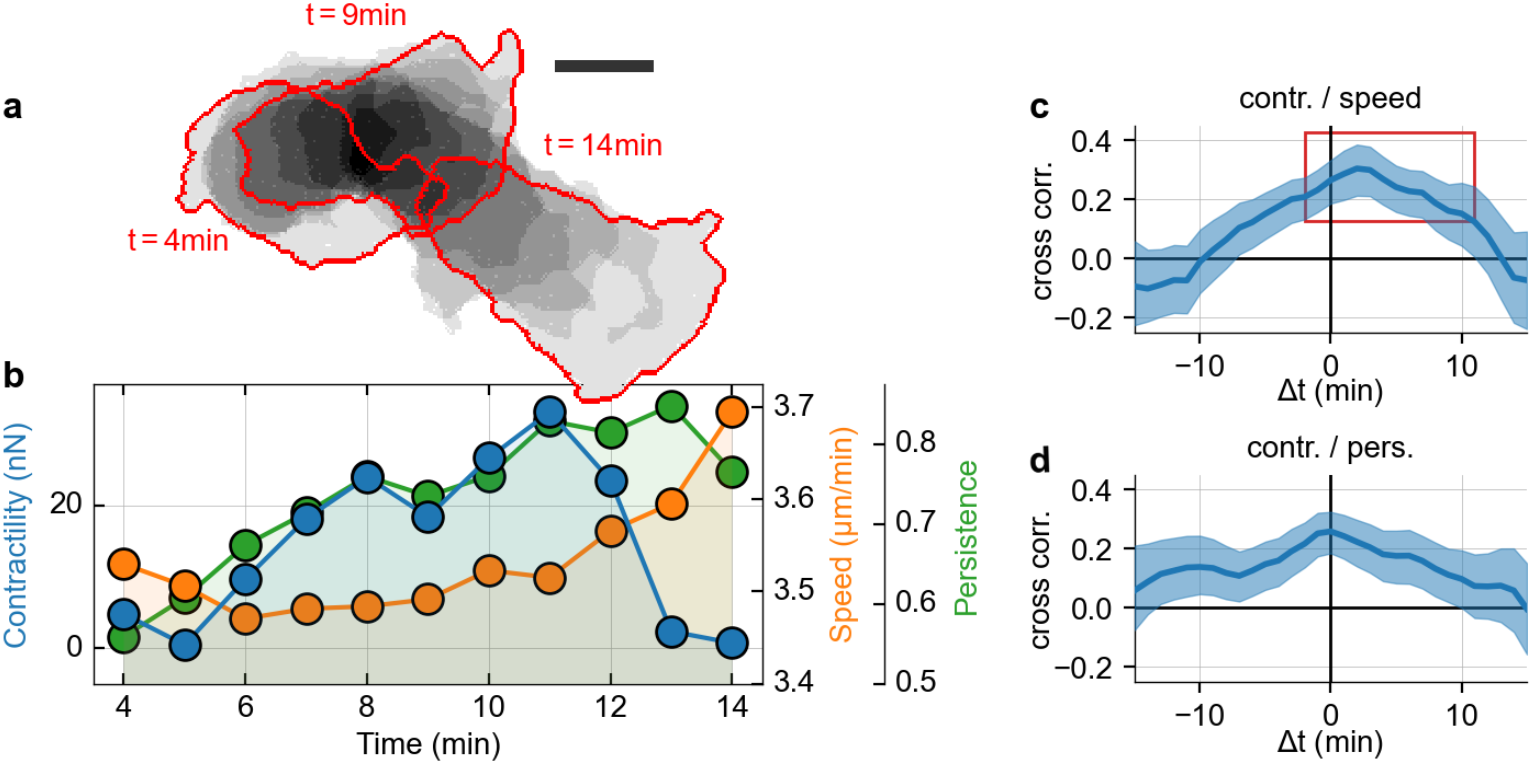
Dynamic variations of contractility during the migration of NK92 cells. **a:** Exemplary migration path of a NK92 cell migrating within a 3D collagen gel for 10min. Cell outlines are shown in red, darker patches indicate a longer retention time at that spot, lighter patches indicate fast changes of cell morphology. Scale bar: 10*µ*m. **b:** Reconstructed cell contractility (blue), cell speed (orange), and directional persistence (green) for the exemplary NK92 cell shown in (a). **c:** Cross-correlation function between cell contractility and cell speed. Positive x-values indicate that speed follows contractility in time, negative x-values indicate that contractility follows cell speed. The correlation function is based on data from n=52 cells and considers all time steps with positive contractility. The red rectangle indicates the right-shifted peak of the correlation function. Error intervals indicate 1sem obtained by bootstrapping. **d:** Same as in (c) but showing the cross-correlation function between contractility and directional persistence.

To identify common patterns in the interplay between cell contractility and cell motility of many individual NK92 cells, we compute cross-correlation functions between contractility and cell speed as well as directional persistence (Fig. 3c,d). Importantly, we find that, on average, cell speed and directional persistence increase with cell contractility, with a small time lag of 2 min, suggesting that NK92 cell motility benefits from contractile forces. This interplay of cell speed, directional persistence, and cell contractility closely resembles the gliding motion during mesenchymal migration of breast cancer cells, but on a faster time scale (cross-correlation functions fall to zero within 10-15 min for NK92 cells, compared to 1 h for MDA-MB-231 cells^8^). This striking similarity in migration dynamics supports our hypothesis that NK92 cells are able to switch to a mesenchymal-like migration mode to maintain their motility in challenging microenvironments.

### Inhibition of acto-myosin-driven forces decreases the motile fraction of NK92 cells

If NK92 cells use traction forces to overcome steric hindrance in 3D collagen gels, we expect that inhibition of cellular force generation will result in a lower motile fraction as more cells become “stuck” in the narrow pores of the gel. To test this hypothesis, we downregulate acto-myosin contraction in NK92 cells by treating the cells with Rho kinase inhibitor or with blebbistatin. We use concentrations that reduce the magnitude (Fig. 4a) and frequency of force peaks (Fig. 4b), but do not completely inhibit cellular force generation so that the cells retain their ability for dynamic shape changes and migration. In agreement with our hypothesis, we find that the fraction of motile NK92 cells is substantially reduced by ∼50% when acto-myosin contraction is reduced (Fig. 4c). Importantly, the treatment does not affect cell viability (Supplementary Figure 5).

**Figure 4:**
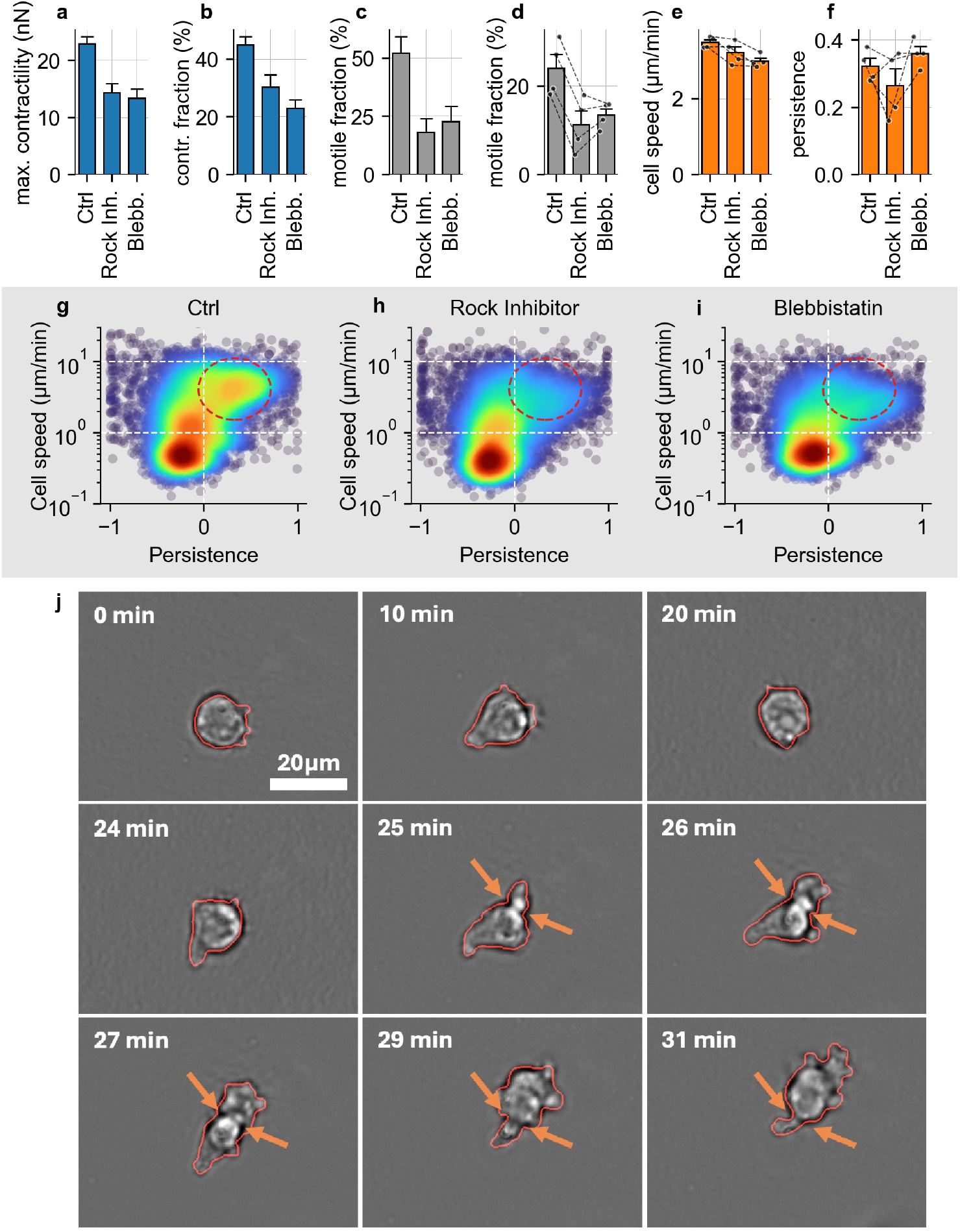
Inhibition of acto-myosin-driven forces in NK92 cells decreases motile fraction, but not motility. **a:** Cell contractility, evaluated by taking the mean over the maximum contractility of each cell during a 24 min observation period (to capture the contractility during the short contractile bursts that NK92 cells exhibit), for the control and for treatment with 10*µ*M Y-27632 Rock-inhibitor and 3 *µ*M blebbistatin in a 1.2 mg/ml collagen gel (Ctrl n=50, blebbistatin n=44, Rock inhibitor n=44 cells) in 3 independent experiments. **b:** Contractile fraction of the NK92 cells, calculated as the average fraction of time steps that the cells reach a contractility >5 nN. **c:** Fraction of motile NK92 cells. A cell is motile if the 5 min bounding-box around its migration path has a mean diagonal length of 6.5*µ*m or greater. **d:** Same as in (c), but measured in the high-throughput 3D migration assay (n=3 independent experiments, n = 1000-3500 cells per measurement and condition, one tailed paired t-test, p<0.0001 for Rock inhibitor, p=0.0051 for blebbistatin). **e:** Mean speed of motile NK92 cells as measured in the high-throughput 3D migration assay, paired for all three conditions. Individual paired measurements are indicated by the black dots and lines (n=4 independent experiments; 1000-3500 cells are measured per measurement and condition). **f:** Same as in (e) but showing the mean directional persistence of motile NK92 cells. **g:** Density plot of cell speed and persistence under control conditions. Each point represents the mean cell speed and persistence of a 5 min trajectory of a single NK92 cell. The color shading represents the Gaussian kernel density estimate (n=4 independent experiments). Red dashed circle approximately indicates the motile fraction of the cell population that migrates at least 6.5 *µ*m from its point of origin within 5 min. **h:** Same as in (g), but for NK92 cells treated with 10 *µ*M Y-27632 Rock-inhibitor. **i:** Same as in (g), but for NK92 cells treated with 3 *µ*M blebbistatin. **j:** Exemplary time-lapse brightfield image sequence of a NK92 cell that remains stuck in the collagen matrix from t=0 min to t=25 min. Note that the cell changes its morphology frequently but is unable to move. After 25 min, the cell appears to protrude into a narrow constriction and becomes migratory again. The red contour shows the cell morphology and the orange arrows show the constriction of the cell due to the collagen pore (scale bar: 20 *µ*m).

To validate this finding, we perform high-throughput 3D migration assays in which NK92 cells are seeded into a collagen matrix and then imaged and tracked at low magnification for 5 minutes in multiple field-of-views. This technique allows us to acquire data on 1000-3500 individual cells per condition and experiment. We perform this additional assay for two reasons: first, there may be a selection bias toward elongated, motile cells in single cell 3D TFM measurements, as the researcher must manually search for and select cells for imaging. Second, the high-throughput 3D migration assay allows us to perform paired experiments for all conditions and obtain data on a much larger population of cells.

The high-throughput migration assay shows a lower motile fraction in all conditions compared to the 3D TFM experiment (Fig. 4d), confirming a selection bias towards motile cells in the single-cell 3D TFM experiment. However, the 50% decrease in the motile fraction after downregulation of acto-myosin contraction remains and is consistently observed in all independent replicates of the experiment (Fig. 4d, g-i). Interestingly, cells that remain motile after treatment suffer only a small ∼10% decrease in cell speed (Fig. 4e), and no change in directional persistence (Fig. 4f).

Acto-myosin activity not only facilitates the exertion of pulling forces on the ECM but also drives cell shape changes via the actin cortex during amoeboid migration. To separate both mechanisms, we place the cells in non-adhesive carbomer gels, thereby forcing them into an amoeboid migration mode. As expected, we find that without adhesion to the ECM, the motile fraction in carbomer is generally lower than in collagen, but we further find that the reduction of acto-myosin contraction after Rho kinase inhibition and blebbistatin treatment has a less pronounced effect of only 20-25% on the motile fraction in carbomer, compared to 50% in collagen (Supplementary Figure 6). This confirms that the decrease of the motile fraction in collagen after partial acto-myosin inhibition is to a large degree attributable to the inhibition of cell-generated traction forces.

Taken together, this result confirms our notion that large cell tractions are critical for overcoming the steric hindrance when encountering narrow pores, but are much less critical for amoeboid motion in less challenging environments. This view is further supported by our observation that cells that become “stuck” are still undergoing rapid morphological changes and continue their migration path once they have succeeded to overcome the obstacle (Fig. 4j, Supplementary Videos 6-8).

### Contractile forces of primary immune cells

The observation of short contractile phases is not limited to the cultured NK92 cell line. Rather, we find that most immune cell types are capable of generating appreciable traction forces. In particular, we find that without expansion, primary human B cells, monocytes, NK cells, T cells, and neutrophils all exhibit contractile phases on the order of ∼5 nN (with occasional peaks around 20 nN) in which they spend 5-10% of the time when placed in a 1.2 mg/ml collagen matrix with a low stiffness of 85 Pa (Fig. 5a, b; Supplementary Fig. 3). We further confirm that traction forces facilitate migration of primary cells, as cell speed and directional persistence increase after force upregulation (Supplementary Figure 7). For all cell types, the fraction of motile cells remains lower compared to NK92 cells and is subject to significant donor-to-donor variation (Fig. 5c).

**Figure 5:**
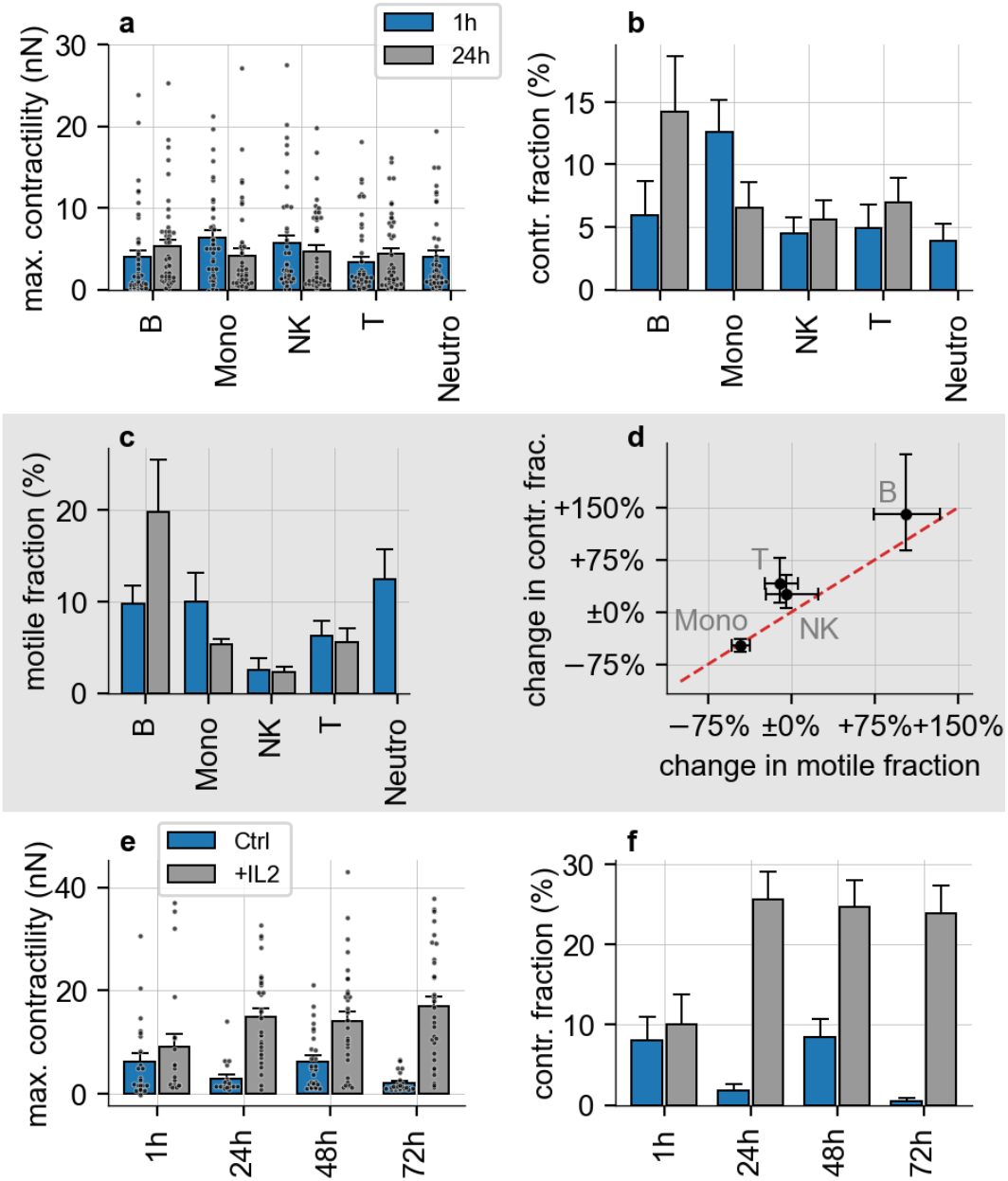
Traction forces of primary immune cells. **a:** Cell contractility, evaluated by taking the mean over the maximum contractility of each cell during the 24min observation period (to capture the contractility during the short contractile bursts) for different primary immune cells 1h (blue) and 24h (gray) after seeding the cells into the 3D collagen gels (B-cells: ‘B’, n=43/44; Monocytes: ‘Mono’, n=39/44; Neutrophils: ‘Neutro’, n=47/0; NK cells: ‘NK’, n=41/43; T cells: ‘T’, n=48/43 in 4 independent experiments). Each black dot represents the maximum contractility of an individual cell. **b:** Contractile fraction of different primary immune cells, calculated as the average fraction of time steps that the cells reach a contractility >5nN. **c:** Fraction of motile primary immune cells as measured in the high-throughput migration assay. **d:** Change in the mean fraction of time steps with a contractility >5nN (called “contractile fraction”) within the first 24h after seeding the cells into the collagen gel, as a function of the change in the motile fraction of the cell population (as measured in the migration assay). The red dashed line indicates the identity line. **e:** Cell contractility of primary NK cells after incubation with IL-2 (gray) and without (control; blue), for different incubation periods prior to seeding the cells into the collagen gel. Each black dot represents the maximum contractility of an individual cell. **f:** Contractile fraction of primary NK cells after incubation with IL-2 (gray) and without (control; blue), for different incubation periods prior to seeding the cells into the collagen gel. Error bars denote 1 SEM in (a)-(c) and (e)-(f), and the 1-sigma credible interval obtained by bootstrapping in (d).

Over a time course of 24 h, B cells increase the frequency of contractile bursts and also increase their motile fraction, whereas monocytes decrease the frequency of contractile bursts and also decrease their motile fraction (Fig. 5b, c). We find a positive correlation between the 24 h change in burst frequency and the change in motile fractions amongst different cell types (Fig. 5d), highlighting the importance of contractile force bursts in the 3D migration process for a diverse set of primary immune cells.

### Migration and contractile forces of ex-vivo expanded NK cells

Ex-vivo expanded NK cells are currently explored for use in adoptive immunotherapy against blood-borne cancers^27,28^. In this type of therapy, patient-derived primary peripheral blood mononuclear cells (PBMCs) are expanded into a population of highly activated NK-cells. Cells are activated by adding IL-2 and culturing them in the presence of IL-15-secreting feeder cells. IL-2 activation is known to promote cell motility^29^, and we reasoned that this effect might be supported by an increased contractility. We confirm that the motile fraction of primary NK cells without IL-2 decreases over a 72 h time period, but remains high when stimulated with IL-2 (Supplementary Figure 8). Similarly, the magnitude and frequency of contractile bursts increase within 1 h after IL-2 addition and remain stable for the following 72 h, whereas contractile forces decline towards zero over this time period in non-stimulated cells (Fig. 5e,f). These data suggest that the enhanced migration of NK cells after IL-2 activation is indeed facilitated by increased cellular force generation.

To relate these findings to a clinical translational setting, we next investigated NK cells that are expanded for 14 days in the presence of high concentrations of IL-2 and IL-15-secreting feeder cells. In general, we find that expanded NK-cells are similarly motile and contractile as IL-2 stimulated primary cells and NK92 cells. However, we find some notable qualitative differences. Specifically, the maximum contractile force and the contractile fraction of expanded NK cells show a bi-phasic response to collagen concentration, reaching the highest values at an intermediate (1.2 mg/ml) collagen concentration (Fig. 6a-d). The motile fraction of ex-vivo expanded NK cells is roughly proportional to the product of the magnitude and frequency of force bursts (c.f. Fig. 2c-e), similar to our findings in NK92 cells. Notably, expanded NK cells retain a higher motile fraction of ∼30% in the high-concentration collagen gel, likely due to their smaller size compared to NK92 cells (Supplementary Figure 9).

**Figure 6:**
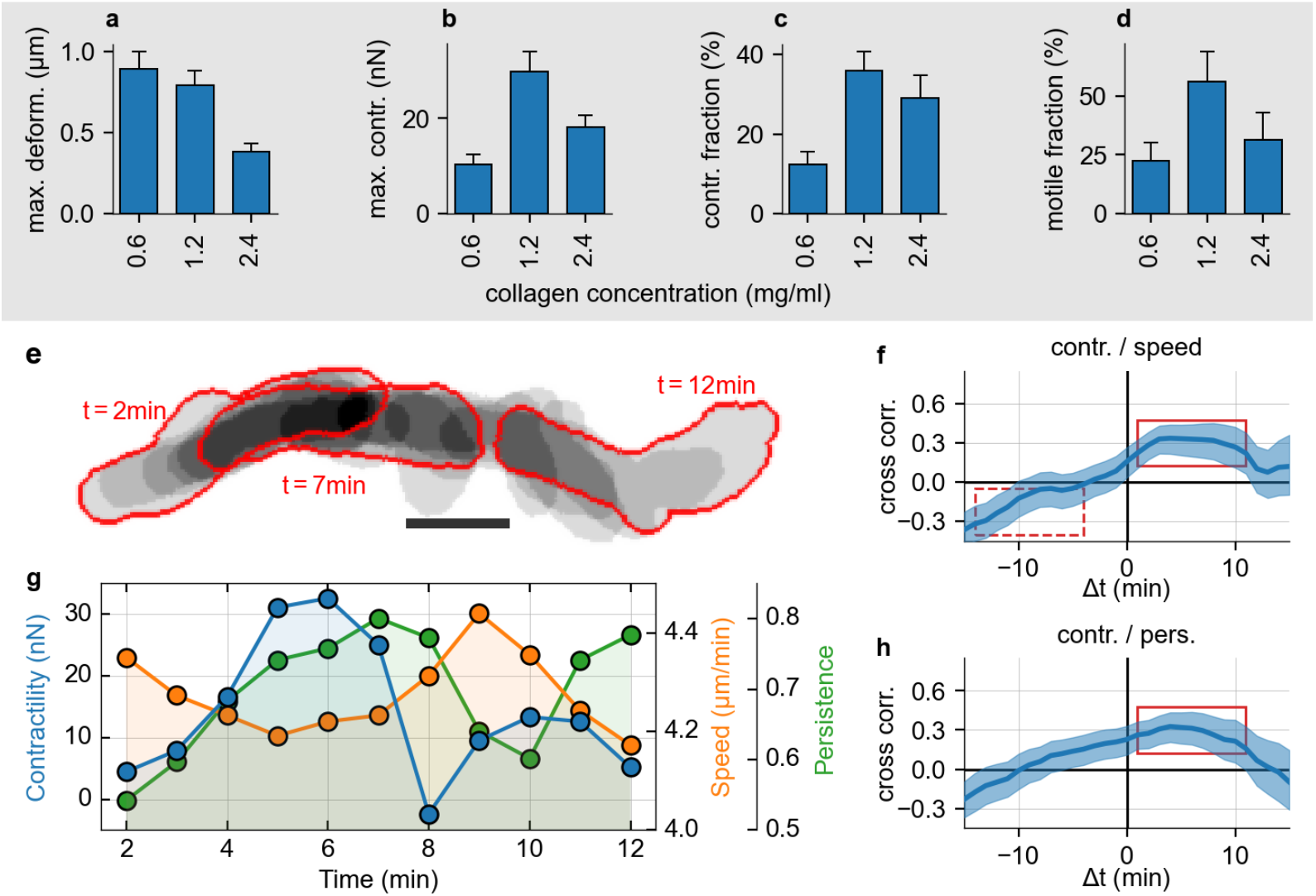
Dynamic variations of contractility during the migration of ex-vivo expanded NK cells. **a:** 99th percentile of the matrix deformations generated by ex-vivo expanded NK cells for different collagen concentrations (0.6mg/ml: n=27; 1.2mg/ml: n=16; 2.4mg/ml: n=16 in 2 independent experiments) **b:** Maximum cell contractility during a 24 min observation period. **c:** Fraction of contractile phases (time periods when cell contractility > 5 nN). **d:** Fraction of motile expanded NK cells for different collagen concentrations. A cell is defined as motile if the 5 min bounding-box around its migration path has a mean diagonal length of 13*µ*m or greater. **e:** Exemplary migration path of an expanded NK cell migrating within a 3D collagen gel for 10min. Cell outlines are shown in red, darker patches indicate a longer retention time at that spot, lighter patches indicate a fast cell movement. Scale bar: 10*µ*m. **f:** Cross-correlation function between cell contractility and cell speed. Positive x-values indicate that speed follows contractility in time, negative x-values indicate that contractility follows cell speed. The correlation function is based on data from n=64 cells and considers all time steps with a positive contractility. The solid red rectangle indicates the right-shifted peak of the correlation function. The dashed red rectangle marks the negative correlation between contractility following cell speed. Error intervals indicate 1sem obtained by bootstrapping. **g:** Reconstructed cell contractility (blue), cell speed (orange), and directional persistence (green) for the exemplary NK cell shown in (e). **h:** Same as in (f), but showing the cross-correlation function between contractility and directional persistence.

In ex-vivo expanded NK cells, we find a more pronounced relationship between traction forces and migration compared to NK92 cells: phases of elevated contractility are followed by phases of elevated speed and persistence lasting for ∼10 min (Fig. 6e-h). Phases of elevated cell speed, by contrast, are followed by phases of reduced contractility lasting for ∼15 min (Fig. 6e-h). Hence, the cell preferentially upregulates traction forces when it becomes slower or less persistent.

### 3D NK cell motility depends on cell-matrix adhesion

The reduced motile fraction of NK cells after force inhibition, and the correlation between contractility and cell speed, provide strong but indirect evidence for our hypothesis that immune cells use traction forces to overcome steric hindrance during 3D migration. Traction forces require strong adhesion to the matrix^30,31^. If NK cells use adhesion-mediated traction forces only when confronted with steric hindrance, we predict that any down-regulation of cell-matrix adhesion will result in a reduced motile fraction (more cells become “stuck”) and a reduced directional persistence (cells take more turns to avoid obstacles). However, we do not expect cell speed to be significantly affected, as motile cells should still be able to migrate in a non-adhesive, amoeboid fashion as long as they do not encounter narrow pores within the matrix.

To test this hypothesis, we modulate cell-matrix adhesion by changing either matrix adhesiveness and/or the ability of the cells to form adhesions. First, we add the adhesion-inhibiting surfactant pluronic to polymerized collagen gels to reduce direct contacts between the cell membrane and individual collagen fibers, forcing the NK cells into a non-adhesive migration mode^32^. For NK92 cells, pluronic treatment reduces the fraction of motile cells by ∼25% and the directional persistence of the remaining motile cells also by ∼25% (Fig. 7a, Supplementary Figure 5). Consistent with our hypothesis, the remaining motile cells maintain their full migratory speed (Fig. 7a). Second, we seed NK92 cells into non-adhesive colloidal carbomer gels (Fig. 7b). These gels are composed of cross-linked polymer particles and provide a soft elasto-plastic microenvironment^21,33,34^ (Supplementary Figure 10, Supplementary Video 9). The relative changes in NK92 cell migration in carbomer gels confirm the results obtained under pluronic treatment, but show a much more pronounced decrease in the motile fraction and directional persistence of motile cells by >75% (Fig. 7b). Strikingly, the migration speed of the remaining motile cells remains unchanged (Fig. 7b).

**Figure 7:**
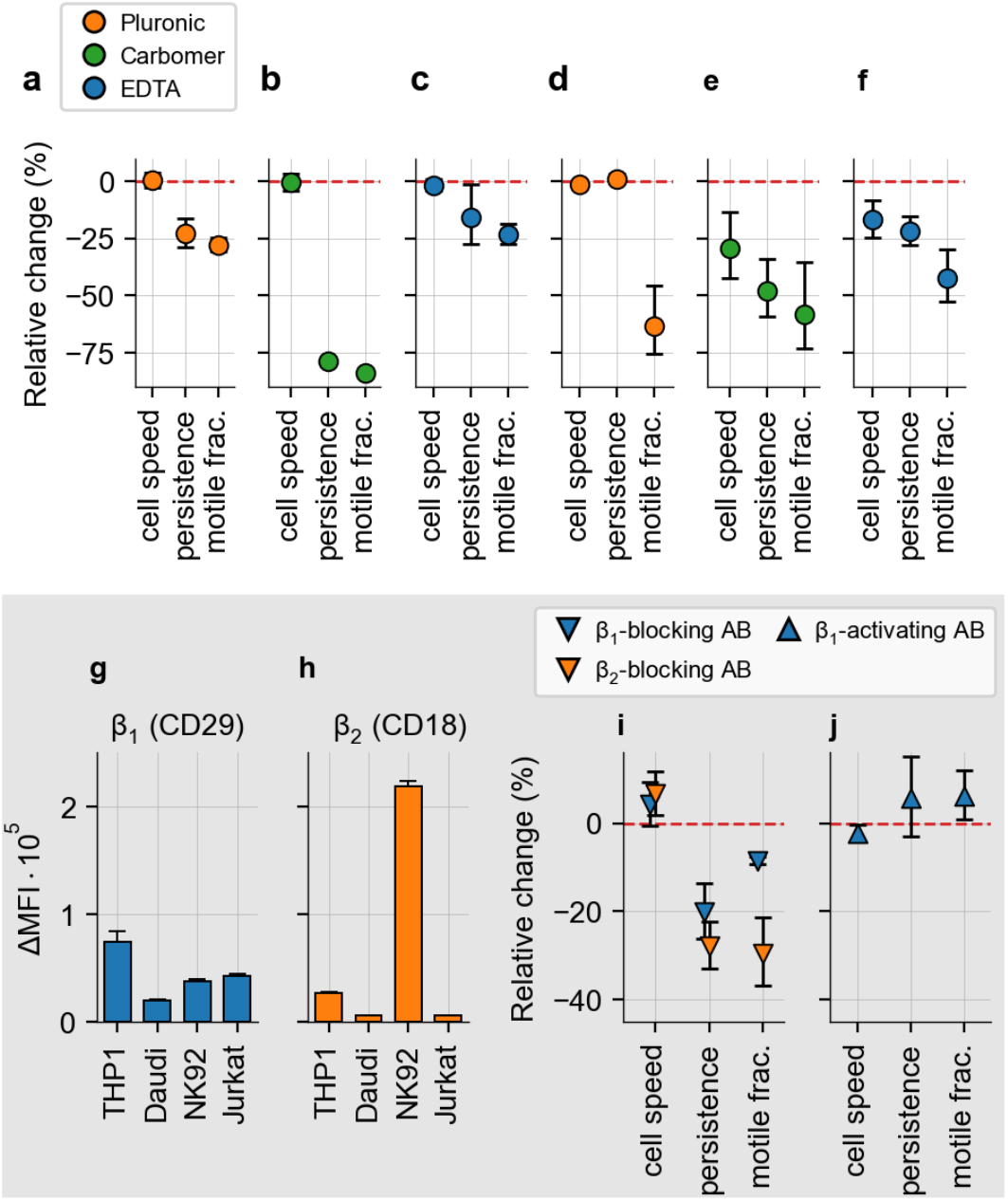
Inhibiting cell-matrix adhesion suppresses NK cell motility. **a:** Relative change of cell speed, persistence, and motile fraction of NK92 cells, measured in high-throughput 3D migration assays (n=3 independent experiments) after inhibiting cell-matrix adhesion externally by the addition of pluronic to collagen gels (hindering cells from making contact with collagen fibers; orange; p=0.02 for the motile fraction, p=0.94 for cell speed, p=0.08 for persistence). **b:** Same as in (a), but for NK92 cells in non-adhesive carbomer gels (n=3; green; p<0.001 for the motile fraction, p=0.90 for cell speed, p=0.001 for persistence). **c:** Same as in (a), but for EDTA-treated NK92 cells in collagen (n=3; blue; p=0.04 for the motile fraction, p=0.56 for cell speed, p=0.38 for persistence), limiting cell-matrix adhesion internally by removing calcium ions that integrins require to maintain adhesion to the matrix. **d:** Same as in (a), but for ex-vivo expanded NK cells (n=2, p=0.24 for the motile fraction, p=0.19 for cell speed, p=0.69 for persistence). **e:** Same as in (b), but for ex-vivo expanded NK cells (n=8, p=0.09 for the motile fraction, p=0.13 for cell speed, p=0.03 for persistence). **f:** Same as in (c), but for ex-vivo expanded NK cells (n=7, p=0.03 for the motile fraction, p=0.11 for cell speed, p=0.03 for persistence). **g:** Delta mean fluorescence intensity of CD29 (β1 integrin) of immune cell lines (THP1, Daudi, NK92 and Jurkat cells), measured by flow cytometry (n=3). **h**: Same as in (g), but for CD18 (β2 integrin) (n=3). **i:** Same as in (a), but for treatment of NK92 cells with a β1 blocking antibody (blue; p=0.01 for the motile fraction, p=0.46 for cell speed, p=0.10 for persistence) and a β2 blocking antibody (orange; p=0.05 for the motile fraction, p=0.26 for cell speed, p=0.02 for persistence) in collagen (n = 3). **j:** Same as in (i), but for β1 (blue) activating antibody treated cells in collagen (n = 3; p=0.37 for the motile fraction, p=0.37 for cell speed, p=0.59 for persistence). Statistical significance is tested using a paired ratio-t test.

We next test how NK cell migration changes when cells are treated with the calcium chelator EDTA. EDTA binds calcium and magnesium ions that are required for the proper function of several adhesion proteins, including integrins^35^. For EDTA-treated NK92 cells in collagen gels, we find that the motile fraction is reduced by ∼25%, the directional persistence of the remaining motile cells is reduced to a lesser extent, and the migration speed of the remaining motile cells is not affected (Fig. 7c). While the effects of pluronic, carbomer, and EDTA on cell migration differ on a quantitative level, they show a similar qualitative pattern: fewer cells remain motile, and the remaining motile cells have to make additional turns within their microenvironment.

For ex-vivo expanded NK cells, the response after manipulation of cell-matrix adhesions is more nuanced. Pluronic treatment results in a large reduction of the motile fraction by >50%, but the remaining motile cells retain their migration speed and directional persistence (Fig. 7d). In carbomer gels and after EDTA treatment, ex-vivo expanded NK cells respond similar to NK92 cells, with a large decrease in motile fraction, a less pronounced decrease in directional persistence, and a smaller decrease in cell speed (Fig. 7e, f). The difference between NK92 cells and ex-vivo expanded NK cells may be attributed to the smaller size of ex-vivo expanded NK cells with respect to pore size and spacing between obstacles in the matrix (Supplementary Fig. 1,7,8).

Pluronic, carbomer, and EDTA broadly inhibit cell adhesion to the matrix without specificity for particular adhesion molecules. Since the dynamic force regulation of NK cells is reminiscent of mesenchymal cells, we further hypothesize that force transmission to the ECM in NK cells is integrin-mediated. To test this hypothesis, we treat NK92 cells with antibodies that specifically block or activate β1 and β2 integrin subunits. Integrin β1 (CD29) is expressed on many cultured immune cell lines, including THP1 monocytic cells,

Daudi B cells, the NK92 cell line, and Jurkat T cells (Fig. 7g). Integrin β2 (CD18) shows a 5-fold higher expression level in NK92 cells compared to integrin β1. The expression of β2 is at least 8-fold higher in NK92 cells compared to the other tested immune cell lines (Fig. 7h). Therefore, we expect that NK92 cells mainly rely on integrin β2 for adhesion^36^, and that blocking integrin β2 will affect the motile fraction more than blocking integrin β1. Indeed, 3D migration assays confirm that blocking integrin β1 decreases the motile fraction of NK92 cells by ∼10%, whereas blocking integrin β2 results in a decrease of ∼30% (Fig. 7i, Supplementary Figure 5). Similar to our experiments in which adhesion is blocked in a nonspecific manner, we again find that directional persistence is significantly lower after integrin blocking. Notably, we find that cell speed is slightly increased after blocking β-integrins, likely because any type of integrin-mediated adhesion slows cells during their default amoeboid migration.

Treatment of NK92 cells with an activating integrin β1 antibody leaves cell motility unchanged across all parameters (Fig. 7j). While we see a small increase in directional persistence and motile fraction, these changes are not statistically significant. This additional finding shows that NK cell motility is not a linear function of integrin activation, but that integrin activity is likely tightly regulated to achieve optimal motility in challenging microenvironments^13^. In summary, manipulation of NK cell adhesion to the ECM demonstrates that NK cells generally achieve rapid migration in 3D biopolymer networks independent of adhesion to the ECM, but are critically dependent on adhesion to maintain motility and directional persistence in the presence of steric hindrance.

### Disruption of the microtubule network increases traction forces and motile fraction

Recent reports have demonstrated that the migration of T cells and their engagement with target cells can be improved by treatment with nocodazole^37,38^. Nocodazole inhibits the polymerization of tubulin and thus destabilizes the microtubule network in cells. Microtubule destabilization has two effects: first, the nuclear deformability is increased^37^, and second, acto-myosin contraction is increased^38,39^. The current understanding is that enhanced immune cell migration under nocodazole treatment is achieved by an increased nuclear deformability as it eases the transition of the cell through narrow pores^2,37^, and by increasing cortical contraction and thus facilitating a fast, contact-guided, amoeboid migration mode in “2.5D” nano-structured surfaces^38^. Here, we provide further context on the effect of nocodazole by investigating cellular force generation and migration of NK92 cells under treatment in 3D collagen gels.

We find that nocodazole treatment substantially increases traction forces of NK92 cells by more than 100% (Fig. 8a). Moreover, treated cells employ traction forces greater than 5 nN more often, in more than 30% of all measured time steps, compared to 15% for untreated cells (Fig. 8b). In accordance with our hypothesis that traction forces help cells to overcome steric hindrance, we find that nocodazole treatment of NK92 cells significantly increases the fraction of motile cells (Fig. 8c). Strikingly, we again see that cell speed of the motile cells is not affected by treatment (Fig. 8d), indicating once more that traction forces do not make cells faster, but instead prevent them from becoming “stuck”. Directional persistence is significantly increased after nocodazole treatment (Fig. 8e), further underlining that higher traction forces alleviate the need to take turns to avoid obstacles within the ECM.

**Figure 8:**
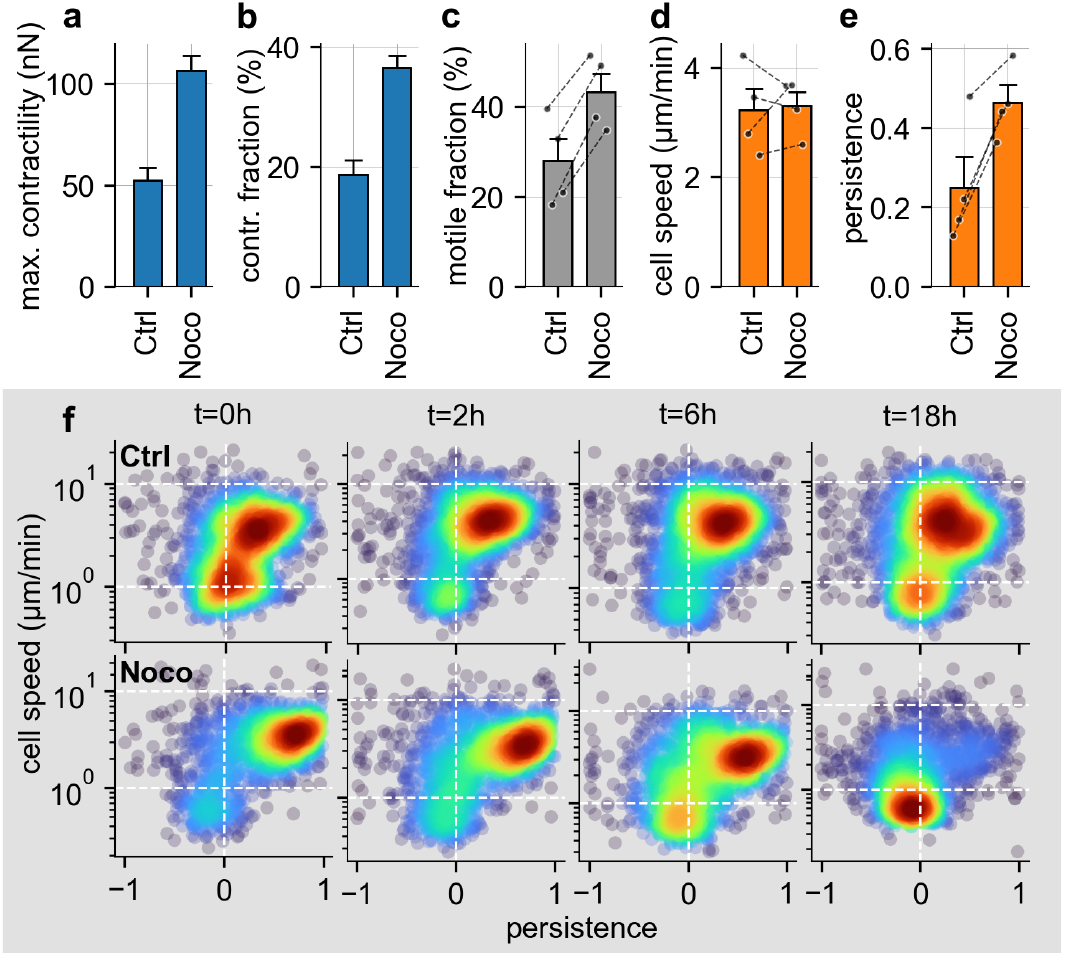
Disruption of microtubule network in NK92 cells increases traction forces and motile fraction. **a:** Maximum cell contractility of NK92 cells during a 24 min observation period for the control and for treatment with 10 *µ*M nocodazole (control: ‘Ctrl’ n=46; nocodazole: ‘Noco’ n=49 cells) in 1.2 mg/ml collagen in 4 independent experiments. **b:** Contractile fraction of the NK92 cells, calculated as the average fraction of time steps that the cells reach a contractility >5 nN. **c:** Fraction of motile NK92 cells measured in the high-throughput 3D migration assay, paired for both conditions. Individual paired measurements are indicated by the black dots and lines (n=4 independent experiments; 3000-4000 cells are measured per measurement and condition, one tailed paired t-test, p=0.001). **d:** Same as in (c) but showing the mean speed of motile NK92 cells (one tailed paired t-test, p=0.409). **e:** Same as in (d) but showing the mean directional persistence of motile NK92 cells (one tailed paired t-test, p=0.005). **f:** Density plot of cell speed and persistence for NK92 cells under control conditions (top row) or for NK92 cells treated with 10 *µ*M nocodazole (bottom row), for different measurement times (0 – 18h) after seeding untreated and treated NK92 cells in collagen. Each point represents the mean cell speed and persistence of a 5 min trajectory of a single NK92 cell. The color shading represents the Gaussian kernel density estimate (n=1 independent experiments, 1500-2000 cells are measured per condition for each measurement time).

Finally, we investigate the effect of an initial nocodazole treatment of NK92 cells on their long-term migration behavior. We find that the initial, strong increase in the motile fraction and in persistence has a short lifetime of < 6 h (Fig. 8f). Importantly, after 18 h measurement time, the majority of all treated cells have become non-motile, whereas the motile fraction of the control cells remains stable (Fig. 8f). The short-term positive effect of nocodazole on NK cell migration thus reverts on longer time scales, leaving most cells in a non-motile state.

## Discussion

We demonstrate that NK cells are capable of exerting significant traction forces on the ECM (in both ex-vivo HAM tissue samples and in-vitro 3D reconstituted collagen networks) that are comparable to the forces exerted by motile mesenchymal cells even after accounting for the smaller size of immune cells. While we confirm the prevailing understanding that immune cells migrate in an amoeboid or friction-mediated manner without significant traction forces^4,12^, we find that immune cells temporarily switch to a highly contractile migratory mode during 10-20% (NK92 cells) and 10-40% (expanded NK cells) of the total measurement time. In particular, NK92 cells show a pronounced mechanosensitivity similar to mesenchymal cells and significantly increase their contractility with increasing matrix stiffness.

Switching between the default, amoeboid migration mode and a more mesenchymal-like migration mode has previously been described in T cells after inhibition of Myosin-IIA, which induces a slower “sliding” migration mode with a continuously progressing contact zone on a 2D substrate^40^. 2D traction forces have been measured in neutrophils^41^ and T cells^42^. Furthermore, leukocytes migrating in dense matrices repolarize into a mesenchymal configuration, placing the nucleus behind the centrosome^43^. Our study complements these findings by measuring pulling forces of immune cells in 3D matrices and, importantly, linking the application of pulling forces to the ability of immune cells to overcome the steric hindrance of a 3D matrix.

To relate cellular force generation to cell motility in 3D, we have shown that both the migration speed and the directional persistence of individual NK cells increase dynamically during and after short contractile bursts. This supports our hypothesis that the switch from an amoeboid to a contractile migration mode is triggered when the cell encounters increased steric hindrance, similar to integrin-driven extravasation of immune cells. This synchronization of cell contractility, persistence, and cell speed is reminiscent of mesenchymal migration of motile cancer cells, which also rely on traction forces to overcome steric hindrance in 3D fiber networks^8,9^.

In support of the hypothesis that immune cells use traction forces to overcome steric hindrance, we further show that inhibition of the acto-myosin contraction in NK92 cells dramatically reduces both contractility and the fraction of motile cells, but has little effect on the migration speed and directional persistence of the remaining motile cells. These results suggest that unless a cell experiences steric hindrance (for example, by migrating through a narrow pore), it retains its default exploratory behavior and amoeboid migration mode. However, with limited acto-myosin contraction, cells are more likely to become “stuck”, resulting in a significantly reduced motile fraction.

The combination of amoeboid migration and occasional contractile bursts is not specific to NK cells. Non-activated primary B, T, and NK cells, as well as monocytes and neutrophils, are all able to exert traction forces on the ECM, although less frequently and not as strong as NK92 cells or expanded ex-vivo NK cells. Interestingly, cellular force generation appears to be closely linked to the adaptation process that immune cells undergo after being seeded into a 3D collagen matrix. Immune cell types that adapt well and increase their motile fraction over 24 hours after seeding also increase the frequency of force bursts, and vice versa. Upon activation with IL-2, primary NK cells increase both the frequency and magnitude of force bursts. Thus, traction force generation may be a key determinant of immune cell infiltration and retention in mechanically challenging environments. Investigating to what extent different immune cell subtypes use traction forces in their respective homing tissues in-vivo would require very stable intravital imaging to detect small matrix deformations in the range of tens of nanometers. This is currently unfeasible, but future improvements may enable a direct verification.

Because large traction forces require a sufficiently strong coupling of the cell’s contractile machinery to the extracellular matrix, it follows that immune cell migration in 3D biopolymer networks depends on cell-matrix adhesion. We have used several methods to inhibit cell adhesion to the ECM. In a non-adhesive environment, which can be achieved either by making collagen fibers non-adhesive using pluronic, or by seeding cells in non-adhesive carbomer gels, we find that the speed of NK92 cells is unchanged, but more cells become “stuck”, and the remaining motile cells take more turns within the matrix. Treating cells with EDTA to weaken adhesion in a non-specific manner confirms these findings. These effects are qualitatively replicated in ex-vivo expanded NK cells. In addition, we have shown that the migration of NK92 cells in 3D fiber networks depends on integrin-mediated adhesion to the matrix. Specifically, we find that blocking integrin β1 and β2 results in a substantial decrease in directional persistence and motile fraction, while cell speed is not reduced. Importantly, the decrease in motile fraction is greater when integrin β2 is blocked compared to β1, consistent with the higher expression level of β2 in NK92 cells.

Finally, we have shown that nocodazole-induced depolymerization of microtubules in NK92 cells results in a substantial, temporary increase in traction forces, motile fraction, and directional persistence, while leaving cell speed of the motile cells unchanged. These results confirm a previously reported increase of the average cell speed under treatment^37,38^, but provide a more nuanced interpretation: not the cell speed of the motile cells changes under treatment, but the fraction of motile cells increases. Hence, nocodazole treatment does not make cells faster but prevents them from getting stuck. Our results suggest that apart from the previously reported increased nuclear deformation^2,37^, traction forces play a crucial role in the nocodazole-induced enhancement of migration.

These in-vitro experiments have implications for current cancer therapies: taxane-based chemotherapies stabilize microtubules to interfere with cell division, but they may further impair tissue– and tumor-infiltration by immune cells^38^. Different chemotherapy agents, such as vinblastine, instead interfere with cell division by destabilizing microtubules, and may thus further enhance immune cell migration^37^, at least on short time-scales. Functional biophysical assays that quantify migration behavior and mechanical interactions between cells and the ECM thus may help to better understand side effects and synergetic effects of existing anti-cancer therapies on immune cell behavior.

In conclusion, this investigation shows that immune cells are able to switch to a mesenchymal-like mode during 3D migration in physiological HAM tissue as well as in reconstituted biopolymer networks. This mesenchymal-like migration mode is characterized by integrin-mediated adhesion to the matrix and large, intermittent traction forces that the cells use to overcome steric hindrance.

## Acknowledgements

This work was supported by the National Institutes of Health Grant HL120839. We acknowledge the support by the Deutsche Forschungsgemeinschaft (DFG, German Research Foundation) project number 540981488 to TC and BF, and project number 460333672 – CRC 1540 Exploring Brain Mechanics (subprojects A01, B01, and C05) to LB, SB and BF. We acknowledge the support by the DFG (FA 336-12/1, TRR-SFB 225) project 326998133 (subprojects A01, B09, and C02) to DB, SB and BF and subproject C05 (TRR225) to RS. We thank Dario Campana, Andreas Baur, and Felix Rico for providing cell lines. This paper was typeset with the bioRxiv word template by @Chrelli: www.github.com/chrelli/bioRxiv-word-template

## Author contributions

C.M., B.F., A.L., C.V. and T.C. designed the study. S.K., G.N., and L.B. performed primary immune cell isolation and FACS analysis. M.W. and C.V. isolated PBMCs from peripheral blood and performed ex-vivo NK cell expansion. PLS and RS isolated and characterized the HAM tissue from human placentas. MWB and MS were responsible for handling of patients and study consent. T.C. and R.G. developed, T.C performed and analyzed short and long-term 3-D migration assays in collagen and carbomer. L.B. and D.B. performed, L.B., D.B., and R.G. analyzed 3-D contraction force microscopy assays. S.B. and D.B. performed collagen and carbomer rheology measurements. E.W. and A.W. developed the neural network-based cell segmentation. T.C., S.K., D.B., L.B. and C.M. performed data analysis. T.C. and C.M. generated the figures. T.C., B.F., C.M., PLS and RS wrote and edited the manuscript.

## Competing interest statement

The authors declare no competing interests.

## Materials and Methods

### Human placenta amniotic membrane isolation

Intact sterile placentae with chorion and amnionic membranes were collected from caesarian sections with informed consent for research use from all mothers. Mothers were seronegative for human immunodeficiency virus (HIV) and human hepatitis B and C. We performed all procedures below under sterile conditions. Whole placentae are washed with 1x PBS and 50 *µ*g/ml penicillin, 50 *µ*g/ml streptomycin, and 1x L-glutamine at least five times to remove remaining blood. The human amniotic membrane tissue (HAM) is then dissected from the chorion layer using sterile forceps and placed in 60 mm tissue culture dishes for further washing with 1x PBS and then RPMI culture media (Thermo-Fisher, USA) containing 10% FCS, 50 *µ*g/ml penicillin, 50 *µ*g/ml streptomycin and 1x L-glutamine (Thermo-Fisher, USA). Phase-contrast microscopy is used to confirm an intact epithelial layer containing polygonal shaped cells as well as for determining the orientation of the tissue. HAM tissue samples are stored for up to 12 days at 37°C, 5% CO_2_ in RPMI culture media that is replenished every day.

### Immunohistochemistry and tissue staining

HAM tissue samples are fixed with formalin and embedded in paraffin, then stained with hematoxylin eosin (Thermo-Fisher, USA) or processed for immunohistochemistry (IHC). The epithelial layer is detected using a mouse anti-human cytokeratin broad spectrum (AE1 & AE3) antibody (1:500, Zytomed Systems, MSK019) and stromal cells within the ECM are detected using a monoclonal mouse anti-human vimentin antibody (clone V9, Sigma-Aldrich, MAB3400). For the detection of primary antibodies, we use the LSAB+ Dako REAL Detection System, HRP (Agilent, Dako) according to the manufacturer’s instructions.

### HAM sample preparation and imaging

To prepare HAM samples for confocal imaging, we carefully cut a 5×5 mm piece using a scalpel and position the ECM-side upwards in the center of a Gene Frame (Thermo Fisher Scientific, cat. no. AB-0576) mounted on a 22×22 mm coverslip (Roth, cat. no. HKE7.1). We then add 20 *µ*l of RPMI containing 120,000 NK92 cells stained for 20 min with calcein (2*µ*M, AAT Bioquest, cat. no. 22004) onto the HAM sample, we seal the frame with a second 22×22 mm coverslip. Every 30 seconds, we image a single z-plane around individual NK92 cells that have invaded into the ECM layer of the HAM sample using confocal microscopy (Leica DM 6000 CFS with HCX APO L 20x/1,00NA W objective). The sample is kept at 37°C during the measurement. CO_2_ content is not controlled as the samples are sealed.

### NK92, Jurkat, Daudi and THP1 cell lines

NK92 cells (purchased from ATCC CRL-2407) are cultured for three weeks prior to measurements in Alpha-MEM medium (Stemcell Technologies) with 15% fetal calf serum (Sigma), 15% horse serum (Gibco), 500 IU/ml human interleukin-2 (IL-2) cytokine (Proleukin S, Aldesleukin, Novartis Pharma, cat. no. 02238131) and 1% penicillin-streptomycin solution (10.000 Units/ml Penicillin, 10.000 *µ*g/ml Streptomycin, Gibco). Jurkat cells (gift from Prof. A. Baur, Department of Molecular Dermatology and Extracellular Vesicle Analysis, Erlangen, Germany), THP1 cells (gift from F. Rico, Aix-Marseille Université, CNRS, Inserm, LAI, Turing center for living systems), and Daudi cells (gift from A. Lux, Department of Genetics, Erlangen, Germany) are cultured for three weeks prior to measurements in RPMI 1640 medium (Gibco) with 10% fetal calf serum, 1% Natrium Pyruvate (Gibco), 1% 100x MEM NEAA (Gibco), 0.3% 1M HEPES (Gibco) and 1% penicillin-streptomycin solution (hereafter called R10 medium).

### Human primary immune cell isolation and storage

Human peripheral blood is collected from healthy donors (Department of Transfusion Medicine and Hemostaseology, University Clinics Erlangen, Germany), with approved consent of all participants and the ethics committee (22-320-Bp) of the university. Immune cells are isolated by density gradient centrifugation using Pancoll centrifugation media (PAN-Biotech, Aidenbach, Germany). Monocytes, NK cells, B cells, and T cells are extracted from the peripheral blood mononuclear cell (PBMC) layer (second layer) with negative magnetic-activated cell sorting (MACS) using the following Biolegend kits: cat. no. 480060 for monocytes, cat. no. 480054 for NK cells, cat. no. 480082 for B cells, cat. no. 480022 for T cells. Granulocytes (mostly neutrophils) are extracted from the bottom layer, whereby the erythrocytes are eliminated by water lysis for 30 s, followed by the addition of 10 vol% of 10x PBS to stop lysis. After cell isolation, monocytes, NK cells, B cells and T cells are stored in R10 medium at 4°C for 24 h, whereas neutrophils are used immediately.

### Flow cytometry analysis

A purity of at least 85% for all isolated primary immune cell subsets (NK, T cells, B cells, monocytes and neutrophils) is confirmed by flow cytometry. Primary immune cells are identified by antibodies against CD45 (Biolegend, cat. no. 368518), a common marker for leukocytes, in combination with a strong forwards and side scatter (for neutrophils) or a second cell-type specific surface marker (CD3 (Biolegend, cat. no. 300408) for T cells, CD19 (Biolegend, cat. no. 302216) for B cells, CD56 (Biolegend, cat. no. 304608) for NK cells and CD33 (Biolegend, cat. no. 303304) for monocytes). Dead cells and cell fragments are identified based on DAPI staining (dilution 1:2,500 of a 5 mg/ml stock solution in PBS with 2% FCS and 0.05% NaN3, AppliChem, Darmstadt, cat. no. 28718-90-3) and excluded from the analysis. Flow cytometry is further used to determine the expression profile of the integrins CD29 (PE anti-human CD29 antibody; Biolegend, cat. no. 303003) and CD18 (PE anti-human CD18 antibody; Biolegend, cat. no. 302107) of the cultured immune cell lines (Fig. 7e – f). All measurements are performed on a FACS Canto II (BD Biosciences), and data are analyzed using FACSDiva and FlowJo Software.

### NK cell IL-2 activation

After 24 h storage at 4°C, primary NK cells are incubated at 37°C in R10 medium with or without 200 IU/ml human IL-2 cytokine for 1h, 24h, 48h or 72h (Fig. 5e – f, Supplementary Figure 8) in a tissue-culture treated 24-well plate (Corning).

### Ex-vivo expansion of NK cells

NK cells are ex-vivo expanded (Figs. 6,7) from PBMCs as previously described^21,44,45^. PBMCs are isolated from healthy donors after oral and written informed consent (Department of Transfusion Medicine, University Hospital Erlangen, Germany; IRB approval number 147_13B and 22-314-Bp), In brief, PBMCs are cultured in the presence of irradiated K562-mbIL15-41BBL feeder cells (gift from Prof. D. Campana, Department of Pediatrics, University Hospital of Singapore) for 14 days in RPMI 1640 medium supplemented with 10% fetal bovine serum, 20 *µ*g/ml gentamycin, 1% L-glutamine and 200 IU/ml human IL-2 cytokine.

### Collagen gel preparation

The collagen solution is prepared at a temperature of 4°C from a 2:1 mixture of rat tail collagen (Collagen R, 2 mg/ml, Matrix Bioscience) and bovine skin collagen (Collagen G, 4 mg/ml, Matrix Bioscience). It is important that all tubes and pipette tips that come in contact with the collagen solution are also cooled to 4°C. We then add 10% (vol/vol) sodium bicarbonate (23 mg/ml) and 10% (vol/vol) 10× cRPMI (Gibco), and dilute the solution to a collagen concentration of 0.6 mg/ml, 1.2 mg/ml or 2.4 mg/ml with a dilution medium containing 1 volume part NaHCO_3_, 1 part 10 × cRPMI and 8 parts H_2_O adjusted to pH 9 using NaOH. Immediately after dilution, the cells are added to the collagen solution, and the cell-collagen mixture is pipetted into 35 mm dishes (ThermoFisher Scientific, Nunc) for traction force measurements, or 6-24 well plates (Corning) for migration measurements. The sample is quickly transferred to the incubator to initiate polymerization. After polymerization at 37 °C, 5% CO_2_ and 95% RH for 1 hour, 1-3 ml cell culture medium or PBS (as described below) is added. Time-lapse imaging of embedded cells is either started directly after this 1h polymerization period (Fig. 5), or 24 h later (for primary immune cells except neutrophils) to ensure that the cells have adapted to the collagen gel and have recovered their full migratory potential.

Two different batches of collagen are used in this work. The rheology of the collagen gels varies between these batches (Supplementary Fig. 1, 3). Batch 1 produced stiffer collagen gels (storage modulus 286 Pa) and is used for measurements shown in Figures 2, 4, 7a-b, 7g-h and Supplementary Figure 11, while batch 2 (shear modulus 84 Pa) is used for measurements shown in Figures 3, 5, 6, 7c-d and Supplementary Figure 8. Rheological measurements are conducted as previously described^18^. In brief, collagens hydrogels (85*µ*l) are polymerized for 30 minutes at 37°C between the plates of a shear rheometer (Discovery HR-3, TA Instruments, Milford, 20mm cone-plate geometry). Subsequently, a frequency sweep (0.2-2Hz) is performed at a low amplitude of 1%, and the stress-strain relationship is measured (0-100% strain) with a constant strain rate of 1%/s. Pore diameters for both batches is 4.4 *µ*m as determined from confocal reflection image stacks (volume of 160×160×200 *µ*m with a voxel sizes of 0.314×0.314×0.642 *µ*m) as described in Refs.^18,46,47^.

### Manipulation of cell-matrix adhesion

Cell adhesion on collagen fibers (Fig. 7a-d) is prevented either by adding pluronic surfactant (1 vol% in PBS, Sigma-Aldrich, cat. no. 9003-11-6), or by calcium ion chelation with 5 mM ethylenediaminetetraacetic acid (EDTA, Roth, cat. no. 8040.3) in PBS. As a control, PBS is used.

Inhibition of Rho-kinase on NK92 cells (Fig. 4) is performed with Y-27632 Rock-inhibitor (Sigma-Aldrich, cat. no. 129830-38-2) dissolved in DMSO (Sigma-Aldrich, cat. no. 67-68-5) at a stock concentration of 1mM. Y-27632 Rock-inhibitor is diluted in cell culture medium to a final concentration of 10 *µ*M. Inhibition of myosin in NK92 cells (Fig. 4) is performed with blebbistatin (Sigma-Aldrich,cat. no. 856925-71-8) dissolved in DMSO at a stock concentration of 10 mM and mixed with cell culture medium to a final concentration of 3 *µ*M. As control, DMSO is mixed in cell culture medium to a final concentration of 0.075 vol%.

Cells are mixed in collagen gel as described above. 2 ml of the PBS-drug solution (pluronic, EDTA, Y-27632, blebbistatin) or 2 ml of PBS with or without DMSO as control are added to the cell samples one hour after initiating collagen polymerization. Cells are incubated for 1 h before measurements.

For integrin blocking or integrin activation with antibodies (Fig. 7g – h, Supplementary Fig. 10), we incubate 5-6*10^5^ NK92 cells prior to mixing them into collagen for one hour in serum-free cell culture medium at 37°C with or without the following antibodies: 10 *µ*g/ml anti-integrin activating β1 (CD29) antibody (Biolegend, cat. no. 303002) and the corresponding IgG1 isotype (Biolegend, cat. no. 400101) as negative control; anti-integrin blocking β1 antibody (P5D2, Abcam, cat. no. ab24693) or anti-integrin blocking β2 (CD18) antibody (Merck, cat. no. CBL158), and the corresponding IgG1 isotype (rndsystems, cat. no. MAB002). After incubation with the antibodies, we mix the NK92 cells in collagen gel as described above.

### Manipulation of microtubules

Destabilization of microtubules in NK92 cells (Fig. 8) is performed with nocodazole (Biomol, Cay13857-10) dissolved in DMSO at a stock concentration of 10 mM. Serum-free cell culture medium is either mixed with nocodazole to a final concentration of 10 *µ*M or mixed with DMSO to a final concentration of 0.1% (DMSO control).

300,000 NK92 cells are incubated for 10 min with serum-free cell culture medium with or without nocodazole. Subsequently, the cells are washed with serum-free cell culture medium and mixed in collagen gel as described above. After one hour of polymerization, 1ml of the serum-free cell culture medium solution with or without nocodazole is added to the gel.

### Carbomer hydrogel preparation

Carbomer hydrogel (Fig. 7a-d, Supplementary Figure 10) is prepared by mixing 9 mg carbomer powder (Ashland 980 carbomer, Covington, USA) with 1 ml R10 medium. The pH is titrated to a value of 7.4 with 10 M NaOH. The carbomer solution is incubated at 37°C and 5% CO_2_ for at least one hour for equilibration. The migration assay is started directly after mixing 300,000 cells in 2 ml of 9 mg/ml carbomer in each well of a tissue-culture treated 6-well plate (Corning).

### High-throughput 3D migration assay

After gel preparation (Table 1) and polymerization, the well plate is transferred to a motorized microscope (Leica DMi6000B, equipped with a 10x 0.25 NA Leica objective and an Infinity III CCD camera, Lumenera) with a stage incubation chamber (Tokai HIT model: WSKMX). For 3D time-lapse imaging, we perform z-scans (10 *µ*m apart) through the 1 mm thick gel every 15 s for a duration of 2.5 min for primary immune cells or for a duration of 5 min for ex-vivo expanded NK cells and NK92 cells. For each condition, we repeat this scanning procedure for 10 fields-of-view in sequence.

**Table 1.** Experimental set-up of the 3D migration assays with different cell types and different collagen concentrations and volumes due to the use of different tissue treated plates shown in several figures.

| Cell type | Cell number | Collagen volume | Collagen concentration | Type of well plate | Figure number |
| --- | --- | --- | --- | --- | --- |
| Non-activated primary immune cells | 75.000 | 450 µl | 0.3, 0.6, 1.2, 2.4 mg/ml | 24 | 5c-d |
| IL-2 activated NK cells | 150.000 | 750 µl | 1.2 mg/ml | 12 | 5e-f, and S4 |
| Ex-vivo expanded NK cells | 300.000 | 1-1.5 ml | 1.2 mg/ml | 6 | 7c-d |
| NK92 cells | 300.000 | 1-1.5 ml | 1.2 mg/ml | 6 | 4d-j; 7a-b; 7g-h; 8; S6, |

For analysis, we detect individual cells using a convolutional neural network that is trained on 4 – 16 manually labeled minimum/maximum intensity projections with approximately 100 cells in each projection. Additionally, we perform data augmentation by image flipping, transposing, and cropping. The network is based on the U-Net architecture^48^. The labeling accuracy of the network ranges from 82% for monocytes to 96% for T cells, 86% for NK92 cells and 94%^21^ for ex-vivo expanded NK cells (F1 score). We then connect the x,y positions of detected immune cells between subsequent images to obtain migration trajectories. An immune cell is classified as motile if it moves away from its starting point by ≥6.5 *µ*m within 2.5 min for primary immune cells, 13 *µ*m within 5 min for ex-vivo expanded NK cells, and 6.5 *µ*m within 5 min for NK92 cells. Cell speed is determined as the diagonal of the bounding box of each cell trajectory, divided by the measurement time of 2.5 min or 5 min. Directional persistence is determined as the average cosine of the turning angles between consecutive cell movements. Zero persistence corresponds to random motion, whereas a persistence of unity corresponds to ballistic motion.

### Single cell 3-D traction force microscopy

3D traction force microscopy is performed as described in Refs.^18,20^. In brief, we pipette 3 ml collagen (0.6 – 2.4 mg/ml) solution with 200,000 cells into a cell culture-treated 35 mm Nunc Petri dish (ThermoFisher Scientific). After incubation for one hour to ensure the complete polymerization of the gel, 2 ml of cell culture medium are added, and the first measurements start immediately. An additional waiting time of 24 hours before imaging ensures that non-activated primary immune cells have properly adapted to the collagen gel. We image z-stacks around single cells with a cubic volume of (123*µ*m)^3^ using a confocal microscope (Leica DM 6000 CFS with HCX APO L 20x/1,00 W objective) equipped with a galvanometric stage and a resonance scanner, which allows for time-lapse imaging of 4 single cells simultaneously with a temporal resolution of 1 stack every 60 seconds. In each independent experiment, up to 12 individual cells are recorded for 24 minutes at 37°C, 5% CO_2_. We select elongated cells for traction force measurements specifically, as these are likely to be motile. From the recorded 3D reflection-channel image stacks, we extract the cell-induced deformations of the collagen fibers using 3D particle image velocimetry as implemented by the SAENOPY Python software package^18^. Based on these 3D deformation fields, SAENOPY computes the contractility of each cell at each measured time step. We exclude deformation fields for which the sum of absolute z-deformations exceeds 45% of the sum of the absolute deformations in all directions, as erroneously large z-deformations can occasionally be caused by the acceleration of the galvo stage. We obtain the cell contractility and force polarity of individual cells from up to 4 independent experiments each.

Data obtained in 3D traction force microscopy experiments shown in Fig. 2g,h, Fig. 3b, Fig. 4a,b, and Supplementary Figures 1 and 3 have previously been reported in the work by Böhringer et al.^18^

### Bayesian filtering of cell trajectories

Bayesian filtering of the individual NK cell trajectories is employed as described in Ref.^26^, but adapted to the fast-moving immune cells. In particular, we use the probabilistic programming framework *bayesloop* to infer the temporal evolution of the filtered cell speed and the filtered directional persistence from a series of (*x*_*t*_, *y*_*t*_) –coordinates that are obtained from the TFM experiments. We ignore the movement of the cells in z-direction as cell movement is more pronounced in the x/y-plane due to the free surface of the collagen gel.

We model the measured cell velocity 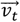 at time step t (with 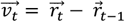, the difference between two subsequent cell positions, 1min apart in the TFM experiments) as an autoregressive process of first order:

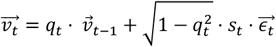

Where *q*_*t*_ denotes the time-varying, filtered directional persistence of the cell, *s*_*t*_ denotes the time-varying, filtered cell speed, and 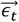 is drawn from a standard normal distribution and represents the driving noise of the stochastic process. Both the filtered directional persistence *q*_*t*_ and the filtered cell speed *s*_*t*_ are allowed to vary slowly over time according to a Gaussian random walk, such that the difference of two subsequent parameter values follows a normal distribution with zero mean and standard deviations σ_*q*_ and σ_*s*_, respectively, so that

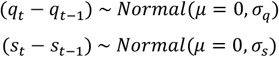

The hyper-parameter σ_*q*_ thus describes how fast the filtered directional persistence *q*_*t*_ changes over longer time scales. Similarly, the hyper-parameter σ_*s*_ thus describes how fast the filtered cell speed *s*_*t*_ changes over longer time scales. In addition to these gradual parameter changes, we account for abrupt jumps in directional persistence and cell speed by assigning a minimal probability *p*_*min*_ to all possible parameter values at each time step. The complete temporal evolution of the filtered directional persistence *q*_*t*_, the filtered cell speed *s*_*t*_, as well as the hyper-parameters σ_*q*_, σ_*s*_, and *p*_*min*_ are then inferred from the series of measured cell positions using the grid-based approach detailed in Ref.^26^ (Supplementary Figure 4).

### Cross-correlation functions

To extract common patterns in the dynamic regulation of cell contractility *c*_*t*_ in relation to cell speed *s*_*t*_ and directional persistence *q*_*t*_, we compute cross-correlation functions *ρ*(*Δt*) across *n* cell trajectories by first creating two vectors *V* and *W* that contain the values (for example of contractility) from all cells pooled together, but shifted in time by the lag-time *Δt*:

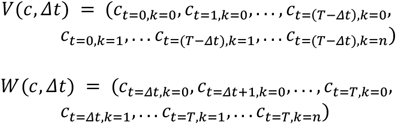

where the index *k* denotes the cell, and the index *t* denoted the time step. Note that we further mask out all time steps for which we have obtained a negative contractility estimate to focus on the dynamics of contractile cellular forces. For positive lag-times *Δt* ≥0, we then compute the cross-correlation function between contractility and cell speed at lag-time *Δt* (i.e. contractility following cell speed with a lag of *Δt*):

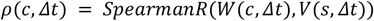

where *SpeqrmanR* denotes the Spearman’s rank correlation coefficient. For negative lag-times *Δt* < 0, we adjust the indexing as follows:

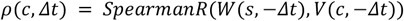

Error bars for cross-correlation functions are determined using bootstrapping, by selecting *n* cell trajectories at random with replacement from the original set of *n* cell trajectories for 1,000 times, resulting in 1,000 bootstrapped correlation values. We subsequently compute the 1-sigma credible interval by taking the 31.73th percentile and the 68.27th percentile as the lower and the upper error bar boundary, respectively.

## Code availability

The 3D traction force microscopy method used in this work is implemented in the Python package *SAENOPY*^18^. The software is open source under the MIT License and is hosted on GitHub (https://github.com/rgerum/saenopy). The Bayesian filtering of cell trajectories is implemented in the Python package *bayesloop*^26^. The software is open source under the MIT License and is hosted on GitHub (https://github.com/christophmark/bayesloop). The detection and tracking of immune cells in the high-throughput migration assays is implemented in custom Python scripts that are available from the corresponding author upon request. The figures in this study have been created using the Python packages matplotlib^49^ and Pylustrator^50^.

## Data availability

Data supporting the findings of this study are available from the corresponding author upon request.

## Supplementary Information

**Supplementary Figure 1:**
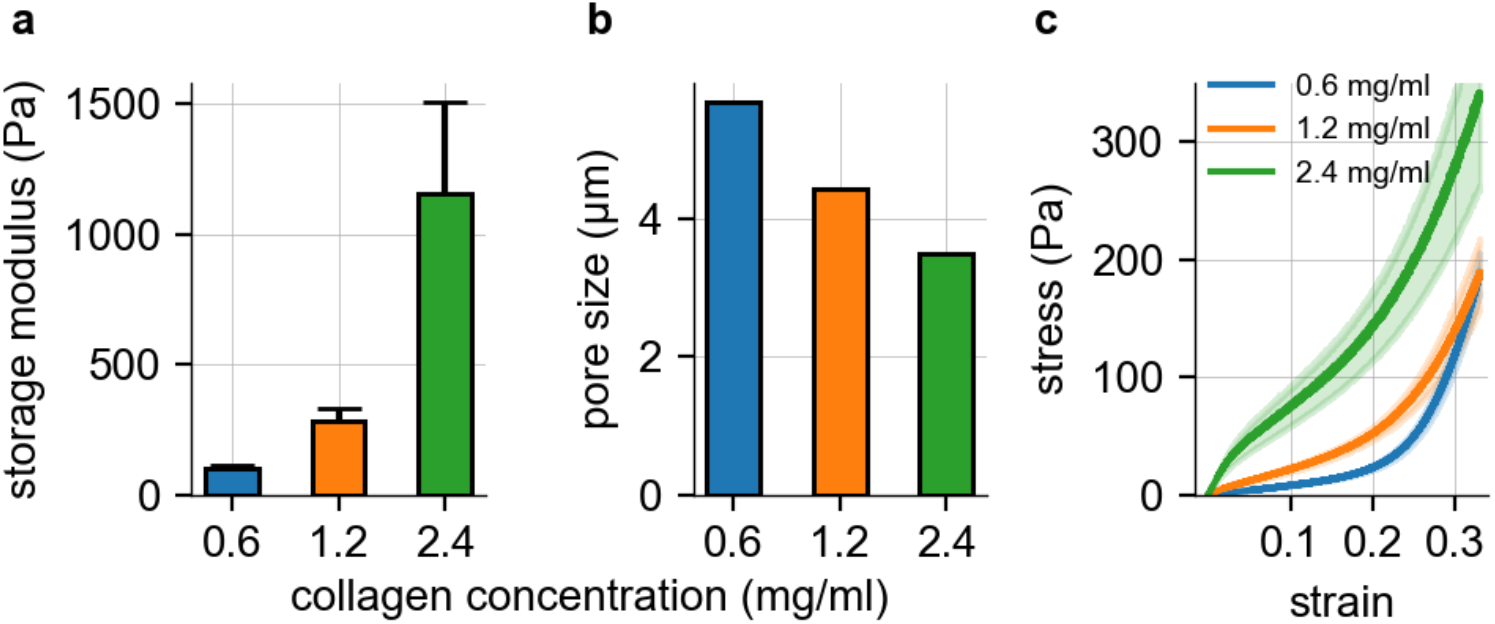
Collagen Rheology and pore size for collagen batch 1. Rheology and pore size measurements of collagen hydrogels at different concentrations of 0.6 mg/ml (blue), 1.2 mg/ml (orange) and 2.4 mg/ml (green) for collagen batch 1 (see Batch C in Böhringer et al.^18^). **a:** Storage modulus scales for small frequency (mean value at 0.02 Hz) for different collagen concentration (0.6 mg/ml: n=3, 1.2 mg/ml: n=4, 2.4 mg/ml: n=4). Stiffness increases with increasing collagen concentration. **b:** Mean pore size for different collagen concentrations averaged from 8 different regions (80×80×100 *µ*m) within an image stack (160×160×200 *µ*m) per concentration (n=1). The pore size decreases with increasing collagen concentration. **c:** Stress-strain relationship of collagen gels for different collagen concentration. Solid lines represent the mean, shaded areas represent the standard deviation (0.6 mg/ml: n=3, 1.2 mg/ml: n=4, 2.4 mg/ml: n=4).

**Supplementary Figure 2:**
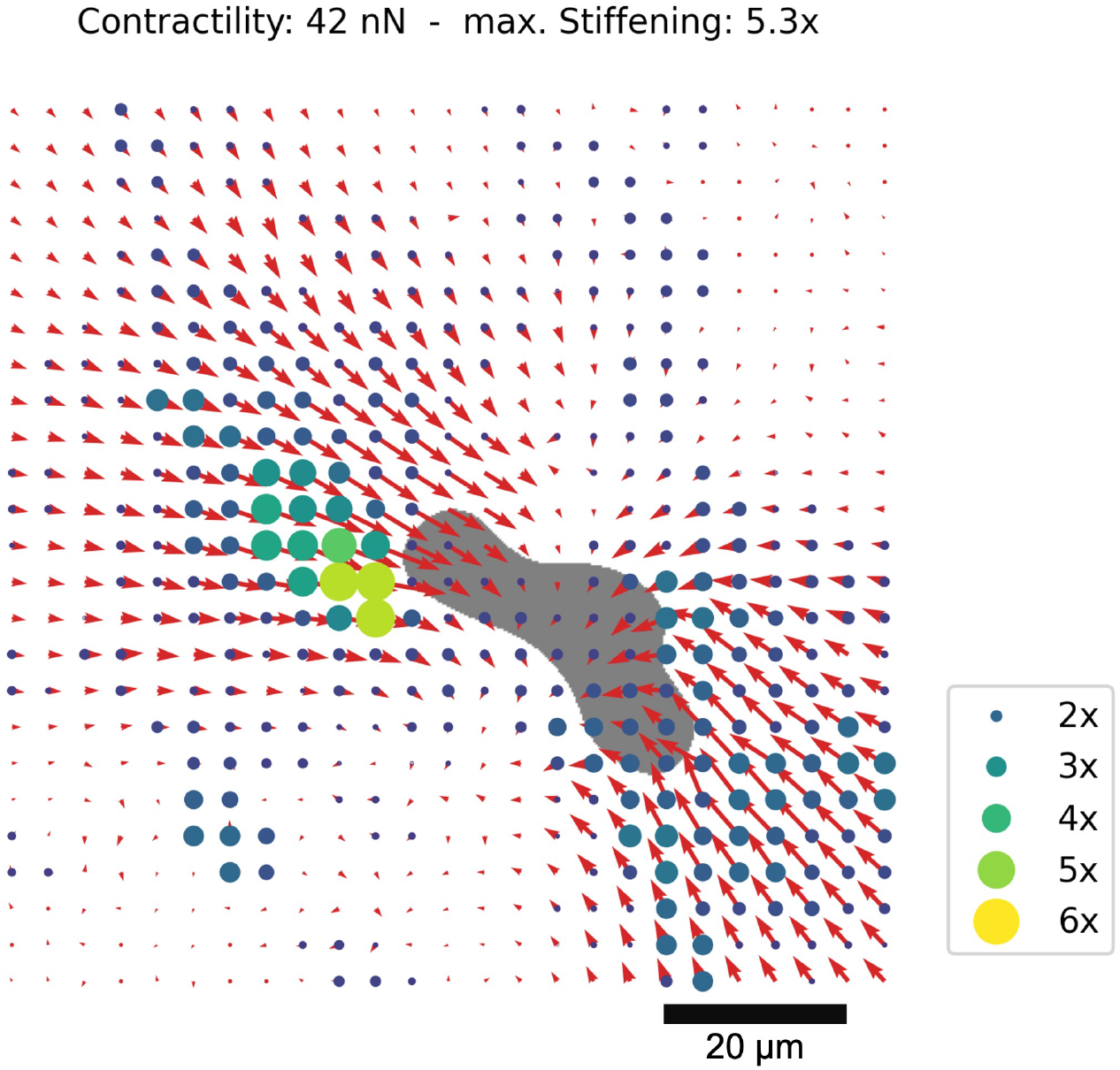
Local collagen strain stiffening by cell-generated traction forces. Exemplary NK92 cell (gray area) migrating in a 1.2 mg/ml collagen matrix (fibers not shown). Red arrows indicate cell-induced matrix deformations (not to scale). Colored disks indicate the local strain stiffening of the matrix due to cell-generated traction forces, as determined by 3D traction force microscopy using a non-linear material model for collagen. Contractility of the cell is 42nN, and results in a local stiffening of the matrix predominantly along the elongation axis of the cell, with a maximal stiffening of 5.3-fold.

**Supplementary Figure 3:**
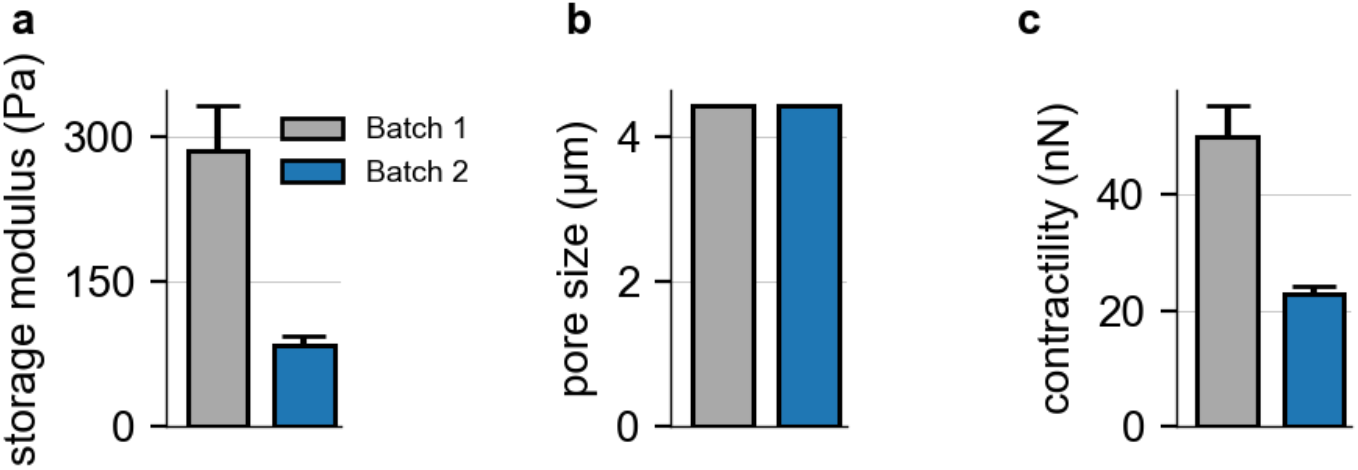
Mechanoresponsiveness of NK92 cells. 1.2 mg/ml collagen hydrogels for two different batches of collagen batch 1 (gray) and 2 (blue) (see Batch A and C in Böhringer et al.^18^). **a:** Storage modulus scales for small frequency (mean value at 0.02 Hz) for batch 1 (n=4) and batch 2 (n=8). The stiffness of batch 1 is more than three times higher compared to batch 2. **b:** Mean pore size measured from an image stack of collagen 1.2mg/ml gels of batch 1 and batch 2. Pore size is averaged from 8 different regions (80×80×100*µ*m) within an individual image stack (160×160×200*µ*m). For both batches, the pore size is 4.4 *µ*m. **c:** Contractility of NK92 cells in batch 1 (n=51) and batch 2 (n=50). The contractility of NK92 cells is twice as high in batch 1 as in batch 2.

**Supplementary Figure 4:**
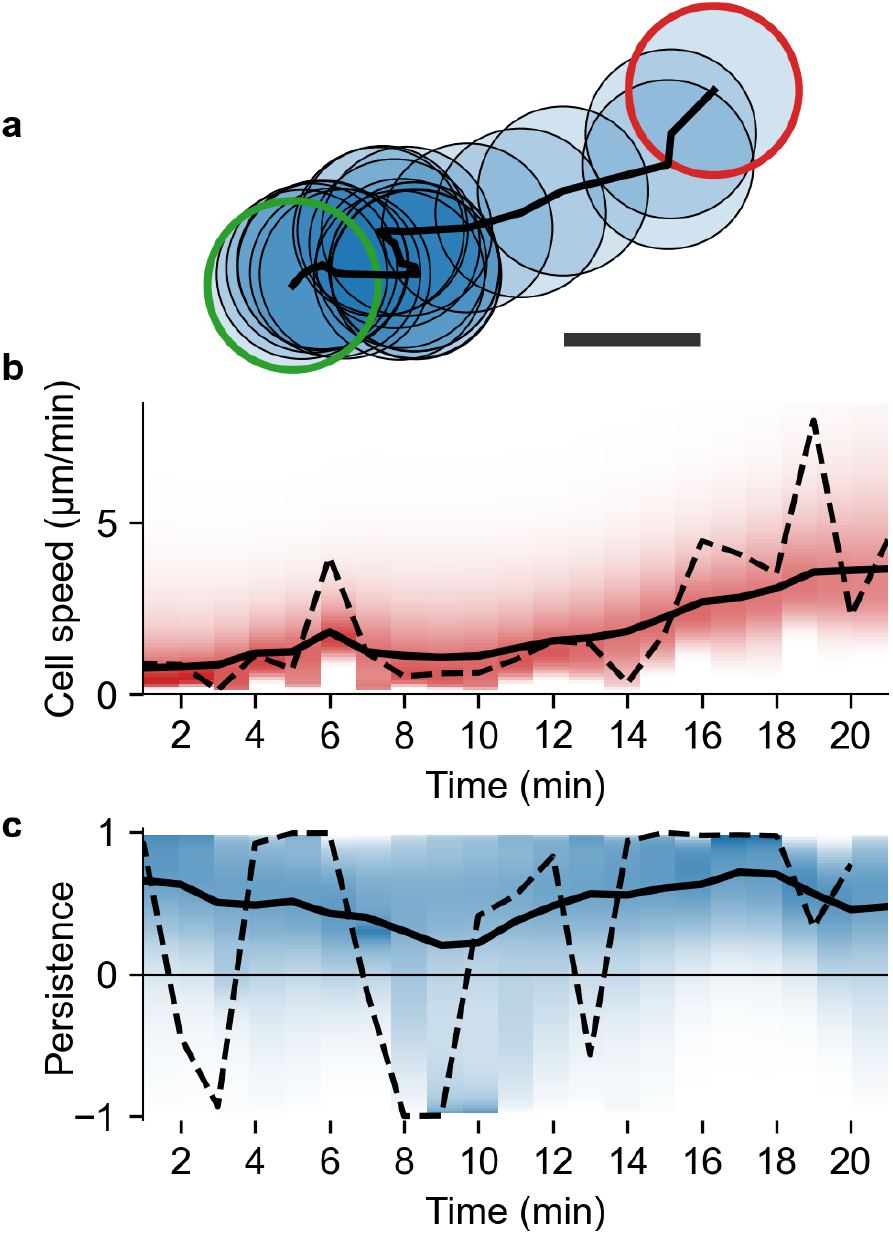
Bayesian filtering of cell speed and persistence. **a:** Exemplary migration path (black) of an NK92 cells embedded in a 1.2 mg/ml collagen gel, measured by confocal microscopy over a time course of 20 min, at an interval of 1 min. The cell trajectory starts at the green circle and ends at the red circle. Scale bar: 10*µ*m. **b:** The momentary cell speed (dashed black line), and the filtered cell speed (solid black line) as obtained from the Bayesian filtering method. The red shading indicates the probability density of the inferred, filtered cell speed. **c:** The cosine of the turning angle as a momentary measure of directional persistence (dashed black line), and the filtered directional persistence (solid black line) as obtained from the Bayesian filtering method. The blue shading indicates the probability density of the inferred, filtered persistence.

**Supplementary Figure 5:**
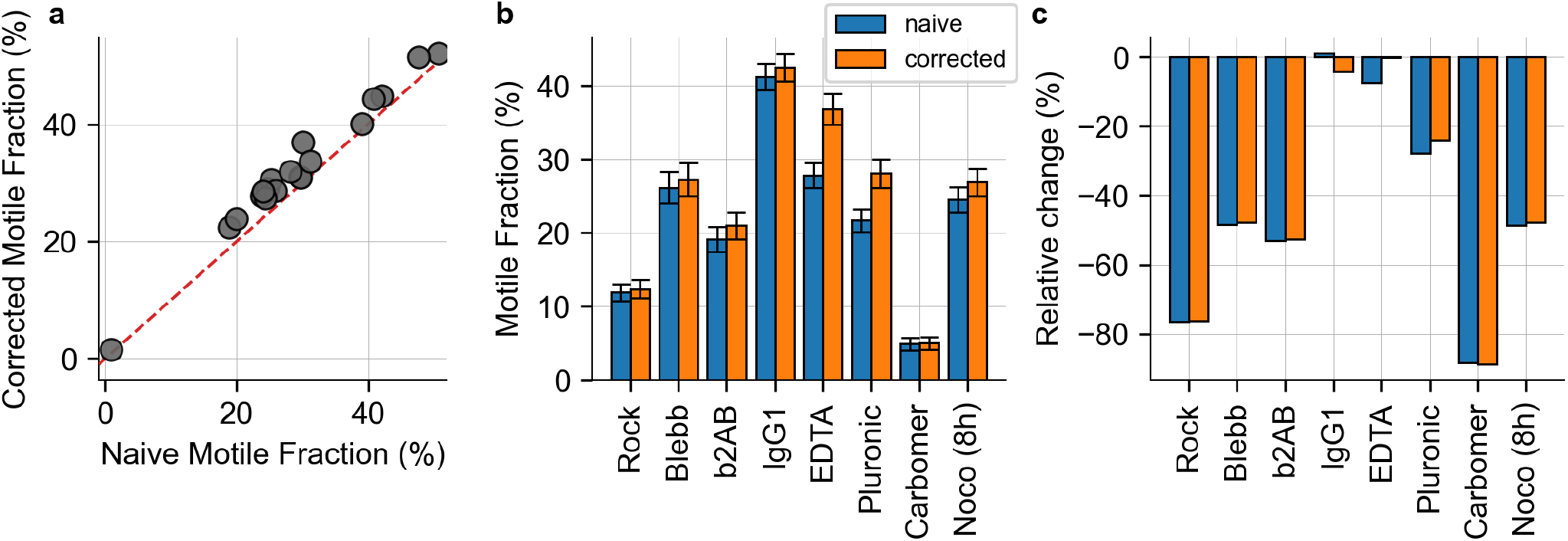
Effect of cell viability on the motile fraction in high-throughput migration assays. To test whether the variations in cell viability due to cell culture conditions and various cell treatments systematically affect the measured motile fraction of cells in high-throughput migration assays, we repeat the migration assays shown in the main text and extend the measurement protocol by a live/dead staining using propidium iodide (Sigma-Aldrich, cat. no. P4864, incubation with 1:1000 dilution of 1 mg/ml for 20 min) at the end of the time-lapse imaging period. By overlaying the detected cell trajectories with the fluorescent images, we determine the number of non-motile dead cells in addition to the number of live motile, and live non-motile cells. This additional metric allows us to compute a corrected motile fraction that is defined as the number of motile cells divided by the number of live non-motile cells. This corrected motile fraction is always equal of greater than the “naïve” motile fraction that does not account for cell viability. **a:** Corrected motile fraction that accounts for cell viability plotted against the naïve motile fraction for n=19 individual experiments using NK92 cells. We do not find any systematic bias of the naïve motile fraction, indicating that variations in cell viability do not systematically affect the outcomes of the high-throughput migration assay with NK92 cells. **b:** Naïve motile fraction (blue) and corrected motile fraction (orange) for different cell treatments (n=1 for each condition; motile fraction of cells treated with Nocodazole is measured 8h after treatment). **c:** Relative change of the motile fraction between control conditions and treatment conditions for the different treatments shown in (b), using the naïve motile fraction (blue) and the corrected motile fraction (orange).

**Supplementary Figure 6:**
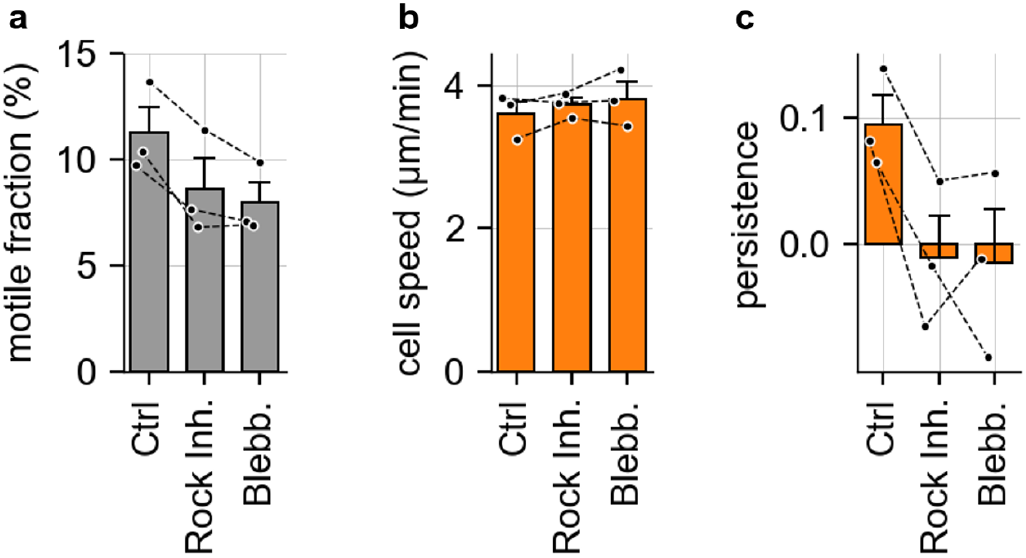
Inhibition of acto-myosin-driven forces in NK92 cells embedded in non-adhesive carbomer gels. Measurement of NK92 cell motility with a high-throughput migration assay, analogous to Figure 4 d-f in the main text, but seeding the cells in non-adhesive carbomer gels instead of collagen hydrogels. **a:** Fraction of motile NK92 cells. A cell is motile if the 5 min bounding-box around its migration path has a mean diagonal length of 6.5*µ*m or greater (n=3 independent experiments, n = 1000-3500 cells per measurement and condition). **b:** Mean speed of motile NK92 cells. **c:** Mean directional persistence of motile NK92 cells.

**Supplementary Figure 7:**
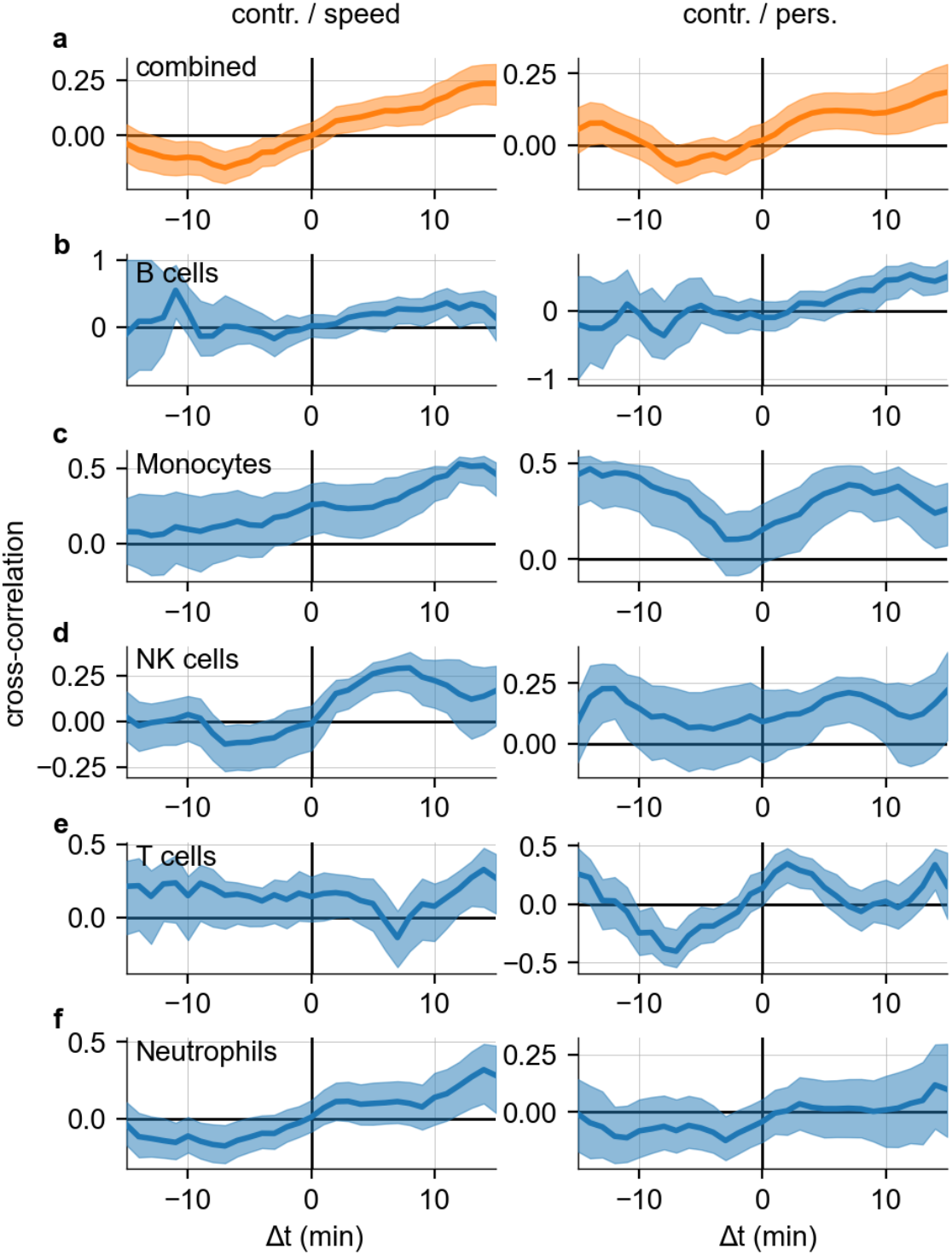
Dynamic regulation of force generation and cell motility in primary immune cells. Left column shows the cross-correlation function between cell contractility and cell speed. Positive x-values indicate that speed follows contractility, negative x-values indicate that contractility follows cell speed. Right column shows the cross-correlation function between cell contractility and directional persistence. Error bars indicate 1 sem obtained by bootstrapping. In general, positive values on the right-hand side of the plots and negative values on the left-hand side of the plots indicate that the cells increase contractility after getting slower or less persistent, and before getting faster or more persistent. **a:** Combined cross-correlation functions computed from all primary cells (n=116). **b:** Cross-correlation functions for B cells (n=20). **c:** Cross-correlation functions for Monocytes (n=28). **d:** Cross-correlation functions for NK cells (n=16). **e:** Cross-correlation functions for T cells (n=16). **f:** Cross-correlation functions for B cells (n=36).

**Supplementary Figure 8:**
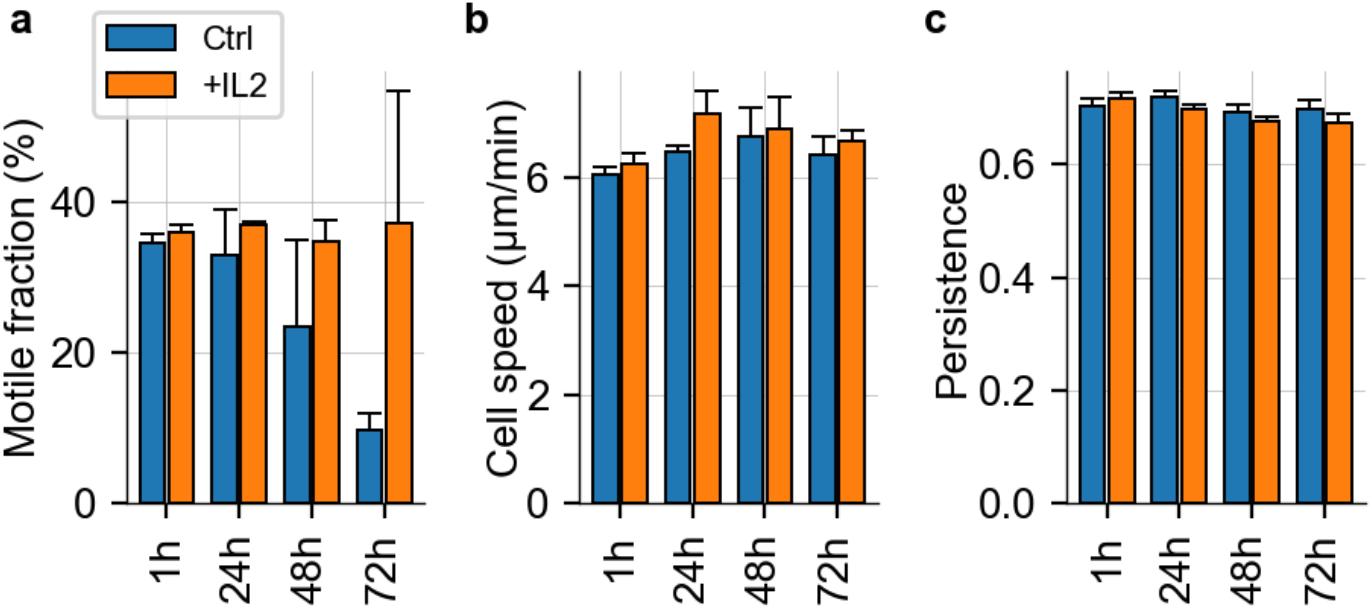
Migration of primary NK cells after IL-2 activation measured in a high-throughput 3D migration assay. Motile fraction, migration speed and migration persistence of primary NK cells incubated with (orange) and without (blue, ctrl) IL-2 from n=2 independent donors for different incubation times (1h – 72h) in 1.2 mg/ml collagen. Each bar represents mean +-se from multiple fields of views (9-20) for each subject. **a:** Motile fraction decreases over time without the addition of IL-2, whereas the motile fraction remains stable over time with the addition of IL-2. **b:** Cell speed of the motile cells increases slightly over time with and without the addition of IL-2, but falls back to the approximate initial state after 72 hours. **c:** Directional persistence of motile cells remains the same over time, with or without the addition of IL-2.

**Supplementary Figure 9:**
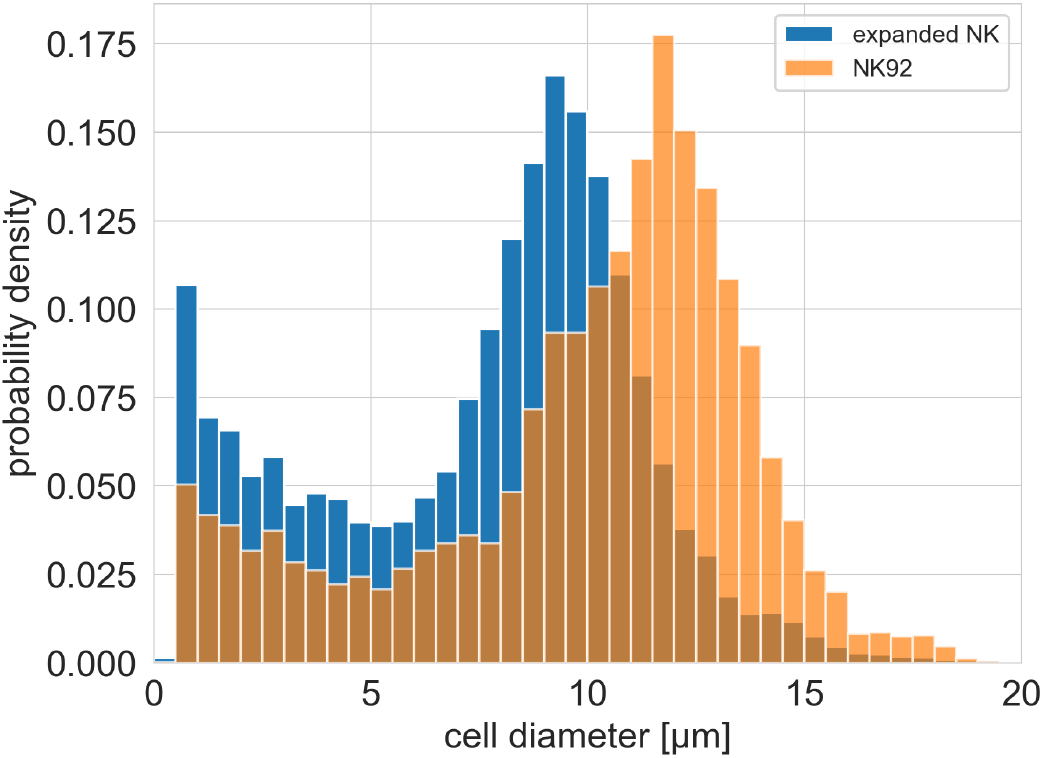
Size of NK92 cells vs. ex-vivo expanded NK cells. Cell size distribution of motile and non-motile ex-vivo expanded NK cells (blue, n = 24.453) and NK92 cells (orange, n = 6.757) using masks generated by the neural network. Ex-vivo expanded NK cells have an average cell diameter of about 7.6 *µ*m, while NK-92 cells have about 9.9 *µ*m.

**Supplementary Figure 10:**
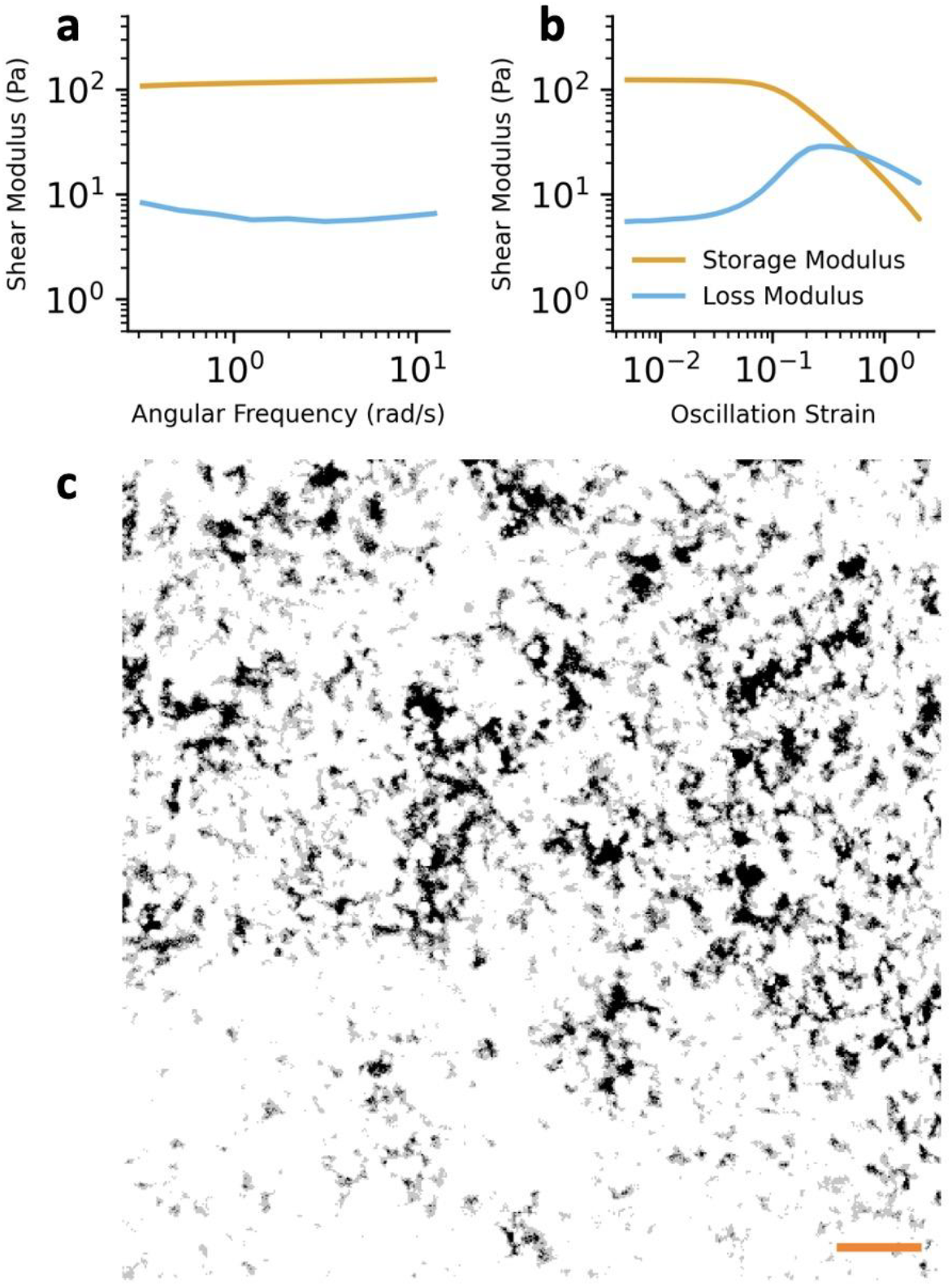
Carbomer rheology and structure. **a:** Shear modulus of Carbomer as a function of angular frequency, measured with a rotational rheometer. **b:** Shear modulus of Carbomer as a function of the oscillation strain, showing the fluidization of Carbomer at high strains. **c:** Patches of low diffusivity (dark) and high diffusivity (light) in Carbomer. Diffusivity was quantified by recording the fluorescent intensity of 1*µ*m-sized beads that are suspended in Carbomer and move randomly due to thermal diffusion (c.f. Supplementary Video 9). Black patches correspond to the 5th percentile of all intensity values that are cumulated over time, and indicate stiff inclusions within the Carbomer matrix. Scale bar is 10*µ*m.

**Supplementary Figure 11:**
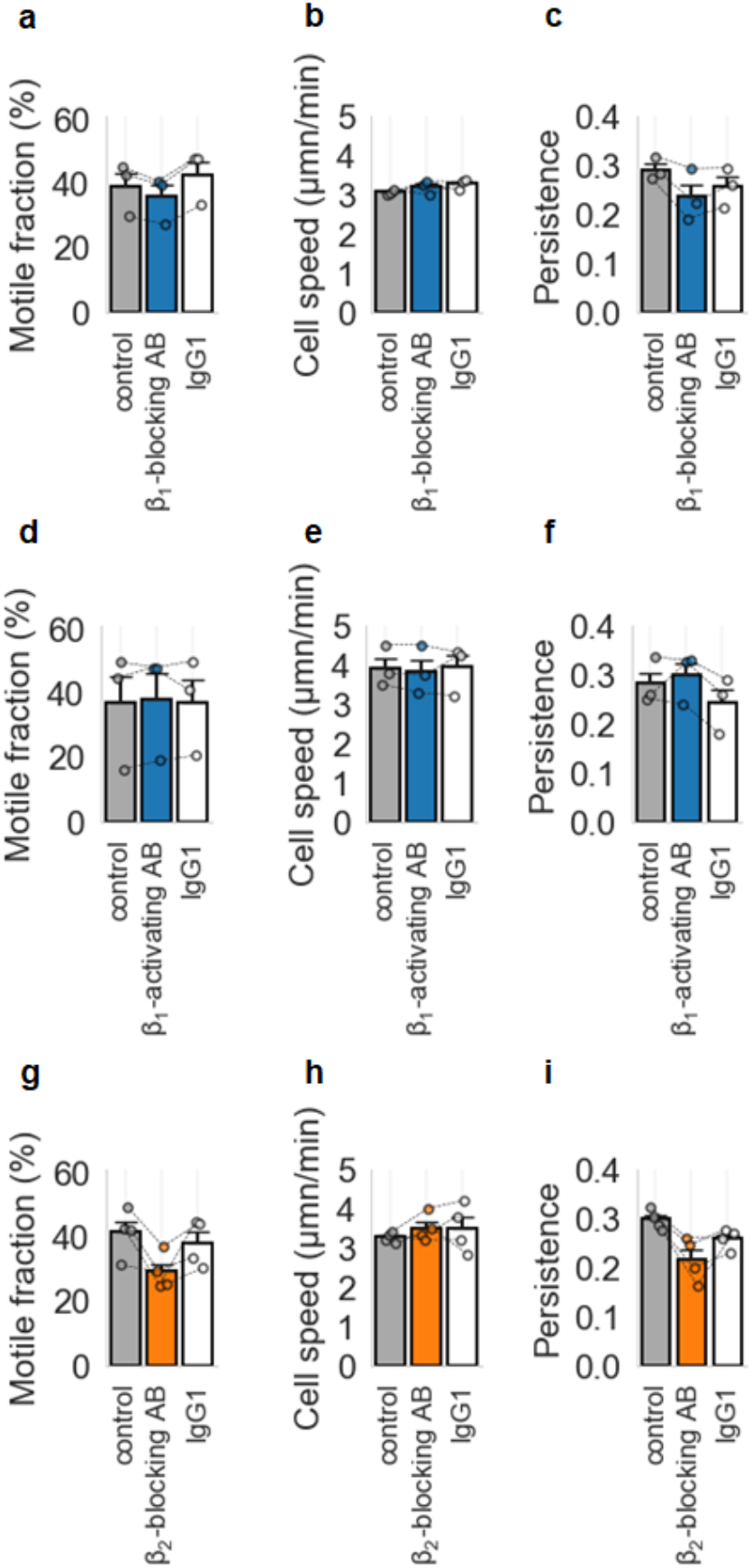
Antibody-based manipulation of cell adhesion measured in a high-throughput 3D migration assay. Motile fraction, migration speed and migration persistence of NK92 cells incubated with blocking (blue or orange) or activating integrin antibody (AB, blue), with the corresponding IgG1 isotype as negative control (white) or without any antibody (gray, control) from n = 3-4 independent samples in 1.2 mg/ml collagen. Each bar represents mean +-se from 10 fields of views for each subject. Each circle represents individual values of each sample. **a:** Motile fraction, **b:** cell speed and **c:** directional persistence of NK92 cells treated with β1 blocking AB (n=3) **d-f:** Same as in A – C, but for NK92 cells treated with β1 activating AB (n=3) **g-i:** Same as in A – C, but for NK92 cells treated with β2 blocking AB (n=4).

**Supplementary Video 1:** Confocal reflection (left) and transmission channel (right) movie of an NK92 cell (red overlay, calcein stain) migrating in the ECM layer of a HAM tissue sample, imaged for 17 minutes and 30 seconds at an interval of 30 seconds. Scale bar: 20*µ*m.

**Supplementary Video 2:** Confocal reflection (left) and transmission channel (right) movie of an NK92 cell (red overlay, calcein stain) migrating in the ECM layer of a HAM tissue sample, imaged for 17 minutes at an interval of 30 seconds. Scale bar: 20*µ*m.

**Supplementary Video 3:** Confocal reflection (left) and transmission channel (right) movie of multiple NK92 cells (red overlay, calcein stain) migrating in the ECM layer of a HAM tissue sample, imaged for 39 minutes at an interval of 30 seconds. Scale bar: 20*µ*m.

**Supplementary Video 4:** Confocal reflection movie of an ex-vivo expanded NK cell (cyan blue, calcein stain) migrating in a 1.2 mg/ml collagen gel (white, confocal reflection channel), imaged for 15 min at an interval of 15 sec. During the first 4 minutes, the NK cell migrates in an amoeboid way without visible deformations of the ECM. After 5 minutes, the cell changes its migration mode by pulling on the collagen fibers. Scale bar: 20*µ*m.

**Supplementary Video 5:** Confocal reflection movie of two ex-vivo expanded NK cells (cyan blue, calcein stain) migrating in a 1.2 mg/ml collagen gel (white, confocal reflection channel), imaged for 15 min at an interval of 15 sec. The cell in the upper right quadrant of the field of view exerts strong pulling forces that result in large deformations of the collagen network. At t=13min, the cell succeeds in pulling its uropod out of the narrow pore and continue its migration path. Scale bar: 20*µ*m.

**Supplementary Video 6:** Long-term time-lapse imaging of NK92 cells migrating in a 1 mm thick 1.2 mg/ml collagen gel for 24 h at an interval of 15 sec (1 sec of the video corresponds to 8 min measurement time). The movie shows the minimum intensity projection of recorded image stacks with a total height of 1 mm at a z-interval of 10 *µ*m.

**Supplementary Video 7:** Brightfield time-lapse imaging at an interval of 15 sec of an NK92 cell migrating through a reconstituted 1.2 mg/ml collagen gel. Initially, the NK cell appears trapped in the dense collagen network. After 25 min, the cell appears to traverse a small constriction and resumes migration. Scale bar: 10 *µ*m.

**Supplementary Video 8** Brightfield time-lapse imaging at an interval of 15 sec of an ex-vivo expanded NK cell migrating through a reconstituted 1.2 mg/ml collagen gel. Initially, the NK cell appears trapped in the dense collagen network. After 31 min, the cell appears to traverse a small constriction and resumes migration. Scale bar: 10 *µ*m.

**Supplementary Video 9:** Confocal images of 0.1 *µ*m fluorescent beads (FluoSpheres carboxylate-modified microspheres 0.1 *µ*m, orange (540/560), ThermoFisher, LOT: 2201623) mixed in 1.5 ml carbomer at an interval of 15 sec. Permanently dark regions in the video indicate stiff inclusions within the carbomer gel, as the beads are unable to diffuse into these regions (some beads become stuck at the boundary of these inclusions). Scale bar: 10 *µ*m.

## Notes

### Competing Interest Statement

The authors have declared no competing interest.

### Summary of Updates

Updated manuscript to include more context on tissue mechanics, and additional results on collagen strain stiffening.

